# De novo designed single-domain antibodies protect against lethal cobra venom neurotoxicity *in vivo*

**DOI:** 10.64898/2026.09.01.748349

**Authors:** Max D. Overath, Emil V. S. Lundquist, Kasper H. Björnsson, Samuel Kerwin, Christian P. Jacobsen, Mark von Bülow Møiniche, Diego Ruiz Espi, Kim Boddum, Melisa Benard-Valle, Monica L. Fernández-Quintero, Stephen P. Mackessy, Timothy P. Jenkins

**Author notes:** Equal contribution (Max D. Overath, Emil V. S. Lundquist).

## Abstract

Generative protein design can now rapidly produce de novo binders with high affinity and functional activity against a wide range of targets, including lethal snake venom toxins. However, so far most reported successes rely on new-to-nature scaffolds with limited therapeutic precedent. Single-domain antibodies (V_H_Hs) offer a clinically validated alternative scaffold that can bind and neutralize long-chain *α*-neurotoxins, which are some of the most lethal components in snake venoms. Here we compare three recently established de novo design models with V_H_H-design capabilities (Germinal, RFantibody, and BoltzGen) for their ability to generate V_H_Hs against the neurotoxin *α*-cobratoxin from the monocled cobra (*Naja kaouthia*). Using standardized model inputs and evaluation criteria based on AlphaFold3 interface confidence (ipTM) and RMSD self-consistency, we find that Germinal was the only method to generate designs passing stringent *in silico* criteria for experimental testing. We therefore performed a larger Germinal design campaign employing three different V_H_H frameworks and experimentally validated 46 designs *in vitro*. Of these, 42 expressed as soluble proteins and we identified four binding hits derived from two of the three tested frameworks. Of the four binders, two lead candidates were further characterized and demonstrated high affinity (K_D_s of 4.1 nM and 10.8 nM), monomeric behavior and low polyreactivity, indicating favorable biophysical and developability properties, as well as functional toxin neutralization *in vitro*. To assess their therapeutic potential we investigated their ability to protect against *α*-cobratoxin toxicity *in vivo*. Both candidates fully protected mice after *α*-cobratoxin challenge, with 100% survival compared to a lethal control. One candidate also retained notable neutralization capacity against whole venom of *Naja kaouthia* with a survival of 56%, while the other protected 22% when tested in a rescue setting. Together, we demonstrate that de novo V_H_H design can generate high affinity single-domain antibodies with *in vivo* protection against lethal cobra venom neurotoxicity, and provide practical insights into method- and framework-dependent performance.

## Introduction

De novo protein binder design, driven by generative deep learning models has rapidly advanced and can routinely produce high-affinity, epitope and target-specific binders without immunization or large-scale library screening [1, 2]. These approaches have shown strong performance across diverse applications, including receptor modulation [3, 4], peptide-MHC binding [5, 6] and the neutralization of lethal snake venom toxins using fully synthetic binders [7].

Despite this progress, the translational path of such de novo binders remains uncertain. Many reported successes rely on new-to-nature protein scaffolds that lack clinical precedent and carry greater uncertainty regarding immunogenicity, pharmacokinetics, and regulatory acceptance compared to traditional biologics [8]. Antibody-based therapeutics represent the largest class of biologics [9] and therefore represent an attractive design scaffold with established clinical precedent, extensive structural and sequence characterization, and widely adopted protocols for translational optimization and therapeutic development [10]. However, while many successful de novo binder design methods rely on rigid *α*-helical or *β*-strand-based interactions, the design of flexible complementarity-determining regions (CDRs) of antibodies that engage a desired epitope remains more challenging.

Fortunately, recent methodological advances have begun to overcome these limitations with several tools now capable of generating epitope-specific antibodies directly from target structures. Notably, the most robust experimental success rate has been reported for single-domain antibodies (V_H_Hs) [11–14]. However, the respective tools enabling the design differ substantially from each other in model architecture, sampling strategies, and filtering criteria, making it difficult to assess practical performance or to identify best practices. Indeed, to date, head-to-head comparisons on shared targets with standardized evaluation metrics remain limited, even at the *in silico* level. Furthermore, experimental validation has so far been restricted to *in vitro* assays, and to our knowledge no V_H_H-based de novo binder has been functionally validated *in vivo* to demonstrate their therapeutic potential.

Snakebite envenoming is a well-suited setting in which to close this gap. This neglected tropical disease causes substantial global morbidity and mortality [15, 16], and recombinant antivenoms based on monoclonal antibodies or V_H_Hs have been proposed as an alternative to plasma-derived products [17–19]. Among the most important targets are long-chain *α*-neurotoxins such as *α*-cobratoxin, which block neuromuscular transmission by binding the nicotinic acetylcholine receptor (nAChR) [20, 21] and are often inadequately neutralized by conventional antivenoms [22, 23]. Generative binder design is well suited to address this problem: since the receptor-binding site of *α*-neurotoxins is structurally well defined, binders can be designed to engage this epitope directly and then assessed for neutralization, rather than recovered through undirected screening. The approach also requires only a target structure, avoiding animal immunization and extensive library screening associated with traditional discovery, and thereby enabling systematic binder design against the large number of toxins relevant to envenoming. While recent work has demonstrated that de novo designed non-antibody binders can neutralize such toxins [7], whether antibody-focused generative design pipelines can achieve comparable performance remains unclear.

Here, we address this gap for V_H_H-based de novo antibody design. We first compared three recently developed V_H_H-capable design methods, Germinal [12], RFdiffusion for antibodies (here referred to as RFantibody) [11], and BoltzGen [13], against *α*-cobratoxin as a medically relevant and structurally well-characterized model target, using a unified AlphaFold3 (AF3)-based scoring and RMSD self-consistency framework to guide method selection. Using the top-performing model (Germinal), we generated and experimentally characterized a panel of designed V_H_Hs, performing comprehensive biophysical, developability, and *in vivo* functional characterization of the most promising binders. This provides an end-to-end demonstration of the translational potential of computationally designed V_H_Hs, showing how de novo antibody design pipelines can generate therapeutically relevant binders.

## Results

### De novo V_H_H design for ***α***-cobratoxin reveals method-dependent design quality

To design V_H_Hs against *α*-cobratoxin, we evaluated three recently developed de novo antibody design models with V_H_H capabilities. To enable a fair comparison, we kept key inputs and evaluation criteria constant across approaches while adhering to each method’s intended design pipeline. Specifically, we used three V_H_H frameworks and constrained each model to design CDRs with lengths matching the corresponding parental framework CDRs (Fig. **1**A). As frameworks, we selected one V_H_H known to bind *α*-cobratoxin (PDB ID: 9GCN [24]) and two V_H_Hs that bind unrelated proteins but have been used successfully as design scaffolds in prior work (PDB ID: 3EAK [25] and PDB ID: 7XL0 [26]). The inclusion of each PDB structure in the training datasets of the utilized methods is summarized in Table S 1. Notably, 9GCN and 7XL0 feature an extended CDR3 conformation, whereas 3EAK has a kinked CDR3 (Fig. S1). For RFantibody and BoltzGen, sequence design was performed using their default inverse-folding modules (ProteinMPNN and BoltzGen-IF, respectively), whereas for Germinal we analyzed designs immediately after the initial generation of a trajectory (prior to AbMPNN refinement). To avoid bias towards Germinal’s internal filters, we included all generated trajectories regardless of their acceptance status in Germinal’s pipeline, providing an unfiltered comparison (cf. Methods).

**Figure 1.**
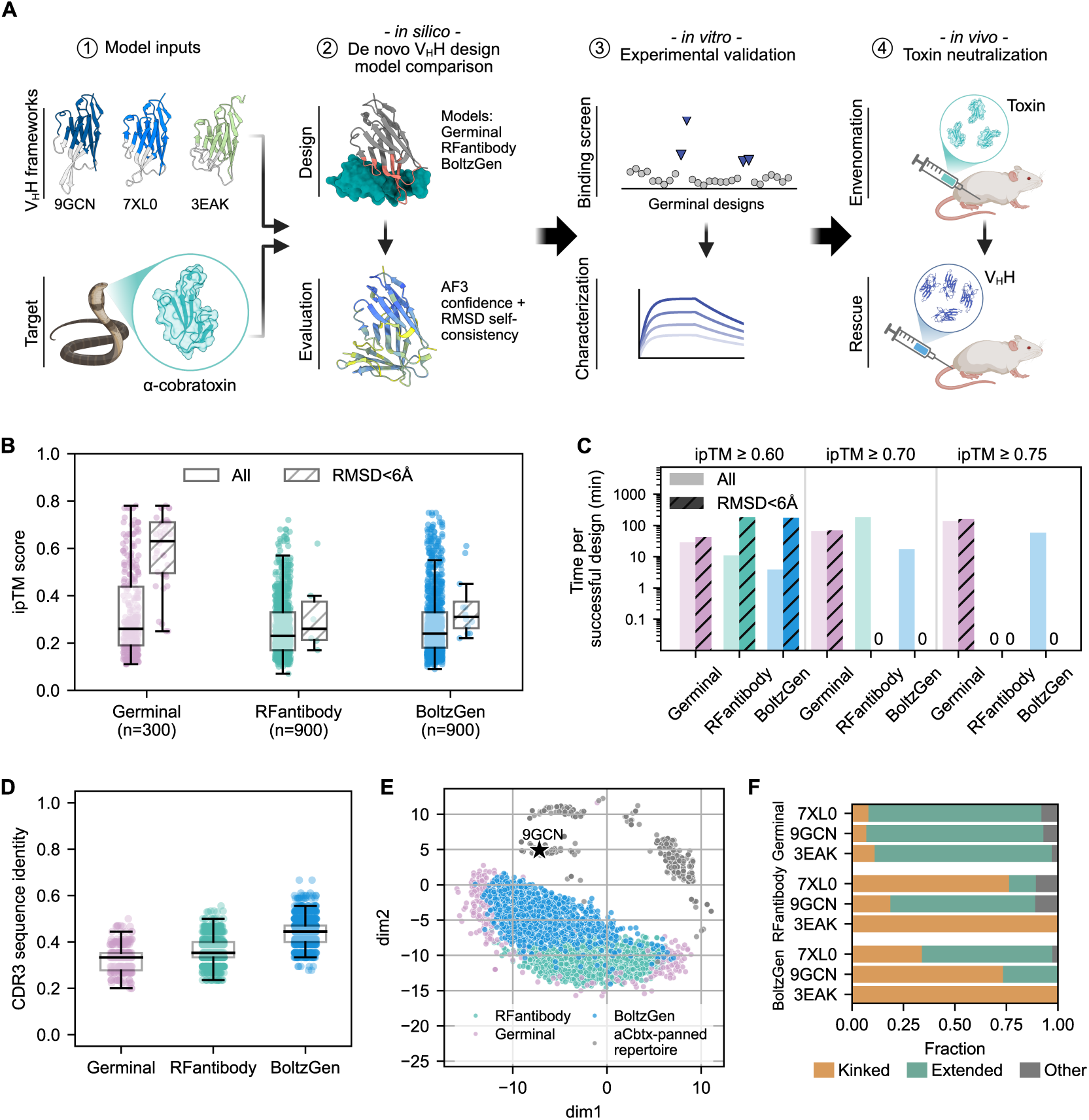
Design approach and model comparison. **A)** Overview of the end-to-end workflow. Three de novo V_H_H design methods were compared in their ability to generate binders against *α*-cobratoxin, employing three different V_H_H frameworks (derived from PDB IDs 9GCN, 7XL0 and 3EAK). Designs were evaluated using AlphaFold3 (AF3) interface confidence and RMSD self-consistency. Based on this evaluation, 46 Germinal designs were experimentally screened for binding. Lead candidates were further characterized for biophysical and developability properties and assessed for their ability to protect against *α*-cobratoxin lethality in an *in vivo* challenge model. **B)** Distribution of interface predicted TM-score (ipTM) for V_H_Hs designed against *α*-cobratoxin, after re-prediction with AF3. Shown are ipTM values for all designs and for the subset retained after filtering for structural self-consistency (V_H_H RMSD *<* 6 Å between the designed model and the AF3 re-predicted structure, computed after superposition of complexes on the target). Here, *n* denotes the number of designs generated by each model prior to RMSD-based filtering. **C)** Compute time per successful design for each method, shown on a logarithmic scale. Success was defined as exceeding an ipTM threshold (0.6, 0.7, or 0.75), shown both with and without the additional RMSD self-consistency filter. A value of 0 indicates that no designs met the corresponding criterion. **D)** Sequence novelty of designed V_H_Hs. Full-length sequences were aligned to the Structural Antibody Database (SAbDab) with MMseqs2; for each design, the closest database match was identified and the corresponding CDR3 sequence identity was reported. **E)** Edit-distance-based multidimensional scaling (MDS) projection of all designed CDR3 sequences compared to naturally occurring CDR3s from an immunized V_H_H library screened against *α*-cobratoxin. 9GCN which was obtained from the same starting library is highlighted. **F)** CDR3 loop conformation of designed V_H_Hs. Fractions of kinked, extended, and other conformations are shown per model and framework across all designs.

To compare designs across models, we predicted V_H_H-toxin complexes for all candidates using AF3 and extracted a unified set of structure- and interface-based metrics. All methods used the same *α*-cobratoxin target structure, and were provided the same set of toxin hotspot residues (positions D27, R33, K35, R36, and V37). Hotspots were selected based on residues contacting the acetylcholine-binding protein (AChBP) in the reference complex and sequence conservation across related *α*-neurotoxins (Fig. S 3 and Methods).

For model comparison, we generated 100 designs per framework with Germinal and 300 designs per framework with BoltzGen and RFantibody, reflecting their higher throughput (Fig. S 2) and aiming for comparable effective sampling depth. All designed V_H_H-toxin complexes were re-predicted with AF3 using a multiple sequence alignment (MSA) for the target only and no MSA for the V_H_H sequence, using one seed per prediction. Designs were first scored by the interface predicted template modeling (ipTM) score, where the median ipTM values were broadly similar across methods (Germinal: 0.26; BoltzGen: 0.24; RFantibody: 0.23), with a modest shift towards higher scores for Germinal (Fig. **1**B). This separation became markedly stronger after applying a filter based on structural self-consistency: we retained only designs with RMSD *<* 6 Å between the designed V_H_H and its AF3 re-prediction after aligning complexes on the target (Fig. **1**B). Such consistency has been proposed as a useful selection criterion [11], and is critical for ensuring that the designed binder maintains the intended binding mode. Under this filter, Germinal produced a substantially larger fraction of high-confidence, structurally consistent candidates than RFantibody and BoltzGen (46/300 designs, median ipTM 0.61, vs. 6/900 and 12/900, median ipTM 0.26 and 0.31, respectively), indicating both higher model agreement and improved interface confidence. A sensitivity analysis across RMSD cutoffs (1–20 Å) confirmed Germinal’s stronger performance on this target at every threshold; the 6 Å threshold was selected as a balance between stringency and ensuring passing designs for all three methods (Fig. S4). To validate these findings with an external, method-agnostic evaluator, we retrospectively repeated the model comparison using ESMFold2 [14] (reported to achieve higher success rates for antibody–antigen docking than AF3, and not used by any of the tested design methods during their own evaluation), and observed a similar performance benefit for Germinal (Fig. S6). We next quantified the required GPU compute time per successful design across different ipTM thresholds (Fig. **1**C). Here, Germinal required considerably more wall-clock time for general design generation than RFantibody and BoltzGen (Fig. S 2); however, when success was defined by exceeding a given ipTM threshold and especially when requiring both ipTM threshold and RMSD *<* 6 Å, the gap in compute time per successful design narrowed substantially. We note that Germinal designs filtered as failed trajectories scored similarly to those with successful trajectories on AF3 ipTM/RMSD self-consistency (Fig. S5A). However, AF3 self-consistency captures only one axis of Germinal’s design requirements: of the 300 evaluated designs, only 70 (23%) completed hallucination with a successful trajectory, and only 17 of these (5.7% overall) passed Germinal’s post-hallucination initial filter (Fig. S5B and Table S2). This indicates that substantially greater sampling is needed to obtain designs meeting Germinal’s complete filtering pipeline.

We next evaluated sequence novelty of all generated designs across the three models by aligning designed V_H_H sequence against the Structural Antibody Database (SAbDab). For each design, we identified the alignment with the highest CDR3 sequence identity as most diversity is normally found in the CDR3. Germinal produced the lowest CDR3 identities overall (median identity =0.33), indicating more novel sequences compared to RFantibody (median identity =0.35) and BoltzGen (median identity =0.44) (Fig. **1**D). This observation is notable given that Germinal uses an IgLM language model to generate antibody-like sequences, whereas RFantibody and BoltzGen relied on general inverse-folding modules for sequence design. Coherent with this finding, Germinal designs obtained a lower hit percentage when aligning against large antibody and V_H_H sequence data bases (Fig. S7). To assess sequence space coverage, we performed multidimensional scaling (MDS) on all generated CDR3 sequences using edit distance. We further compared the designed sequences to naturally occurring CDR3s from an immunized V_H_H library screened against *α*-cobratoxin [27], which served as a reference for the natural sequence landscape (Fig. **1**E). The three methods occupied largely distinct regions, with Germinal forming two well-separated clusters, indicating broader exploration of CDR3 sequence space. Notably, the immunization-derived *α*-cobratoxin-binding CDR3s were distant from all model-generated sequences. We also show that high-confidence designs are not restricted to defined regions of the sampled CDR3 sequence space (Fig. S8). Finally, we analyzed CDR3 conformations using a previously proposed geometric classification into kinked, extended, and other states based on established backbone angle descriptors (Fig. **1**F; [28]). Germinal predominantly generated extended CDR3 conformations across all three frameworks. In contrast, RFantibody produced mainly kinked CDR3s for 7XL0 and 3EAK, but predominantly extended CDR3s for 9GCN. BoltzGen generated mostly kinked conformations for 9GCN and 3EAK, while producing a higher fraction of extended conformations for 7XL0.

### Large-scale Germinal campaign yields experimentally binding V_H_Hs

Since Germinal produced the most promising candidates in our head-to-head model comparison, we performed a larger-scale Germinal design campaign using the three selected V_H_H frameworks and the predefined *α*-cobratoxin hotspot residues, with the goal of experimentally validating a subset of these for binding. We generated roughly 8000 designs in total, of which *∼* 3% passed Germinal’s internal filters (see Methods for applied filters). From these predictions, we down-selected 48 designs across the three frameworks (3EAK: 8; 9GCN: 20; 7XL0: 20). The 48 selected V_H_Hs were then produced recombinantly in *E. coli* (SHuffle T7) and purified via C-tag affinity chromatography. Of the 48 designs, two (both 7XL0) failed during cloning; of the remaining 46, protein concentrations after purification were quantified to assess expression success. In total, 42/46 designs yielded measurable protein concentration, including 8/8 from the 3EAK framework, 19/20 from 9GCN, and 15/18 from 7XL0 (Fig. **2**A). This high overall expression success rate (*∼* 91%, 42/46) is consistent with Germinal’s stringent filtering steps for liabilities such as spatial aggregation propensity (SAP) of the CDRs [12]. All purified candidates were then screened for binding by flow-induced dispersion analysis (FIDA) [29] using AF647-labeled *α*-cobratoxin. Binding was assessed by monitoring the hydrodynamic radius (*R_h_*) of AF647-labeled toxin, where an increase in *R_h_* indicates complex formation. We identified four candidates with a clear increase in Δ*R_h_* between toxin alone and toxin incubated with V_H_H: two from the 9GCN framework (D2, E2) and two from 7XL0 (G3, H3) (Fig. **2**B), with an overall success rate of 8.7% (4/46). Initial one-point BLI screening identified D2 and E2 as the strongest binders, with substantially higher response than G3 and H3 (Fig. S9).

**Figure 2.**
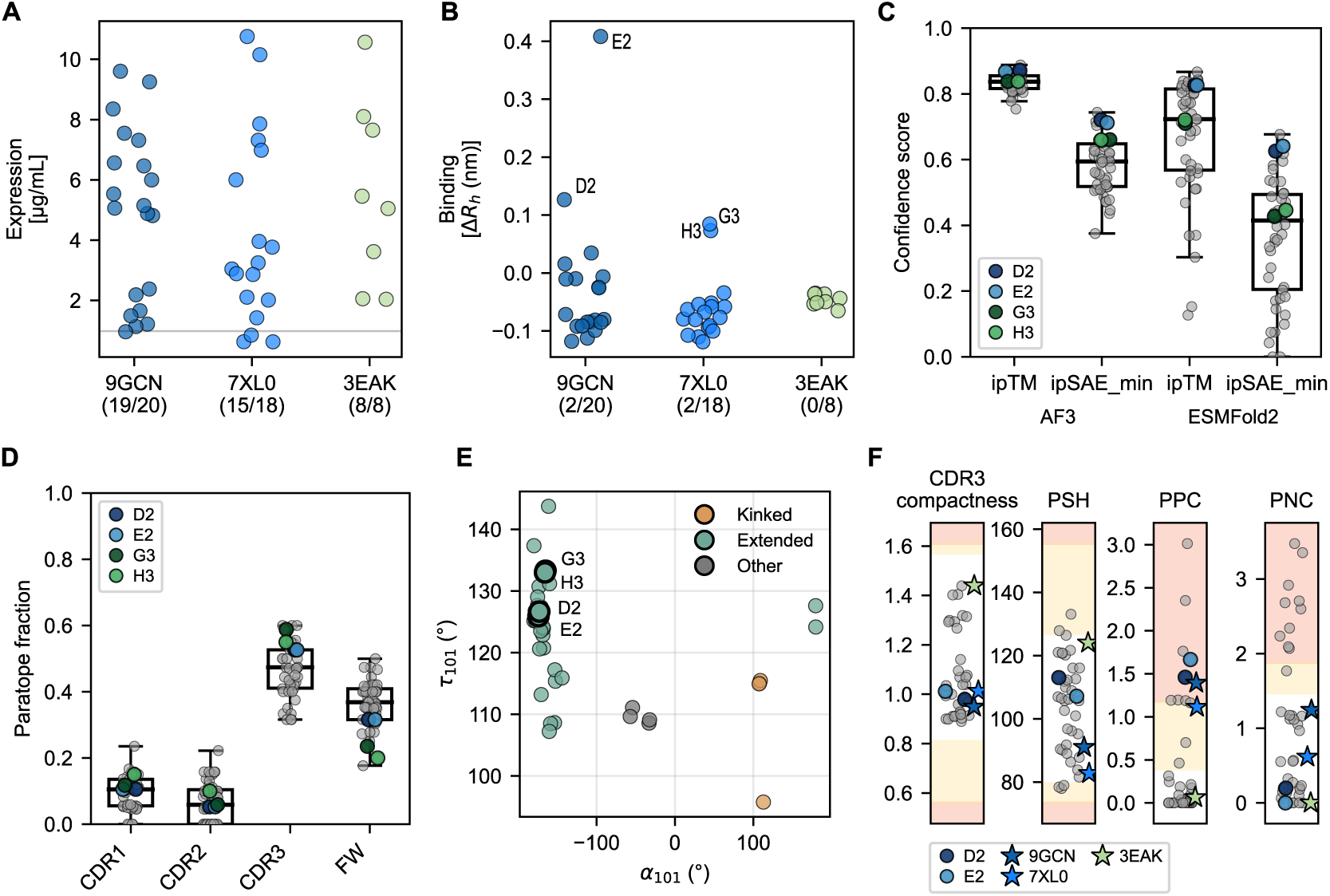
Experimental screening and *in silico* evaluation of 46 Germinal-designed V_H_Hs. **A)** Expression yields after purification (as *µ*g per mL culture volume), shown by framework. The number of designs exceeding the expression threshold is annotated for each framework; the horizontal line indicates the negative-control background used to define the threshold. **B)** Binding screen by FIDA. For each V_H_H, the Δ in hydrodynamic radius (*R_h_*) is shown relative to AF647-labeled *α*-cobratoxin alone and after incubation with the purified V_H_H. Counts of identified hits are annotated per framework. **C)** Retrospective evaluation of ranking true binders based on AF3 as well as ESMFold2 ipTM and ipSAE min scores. **D)** Contribution of CDR1, CDR2, CDR3, and framework residues to the AF3 predicted paratope, reported as the fraction of paratope residues falling in each region. Highlighted are the identified hits. **E)** CDR3 conformation of selected V_H_H based on *α*101 and *τ* 101 angles. All confirmed hits display an extended CDR3. **F)** Developability profiling of the 46 selected designs and the three frameworks with the Therapeutic Nanobody Profiler (TNP), shown as metric values across four CDR-vicinity liability metrics: CDR3 compactness; PSH, CDR-vicinity patch of surface hydrophobicity; PPC, CDR-vicinity patch of positive charge; PNC, CDR-vicinity patch of negative charge. Shaded bands mark the amber (outer 5% of the clinical distribution) and red (outside the observed clinical range) liability regions, with the unshaded region corresponding to the clinical-stage reference range. The three frameworks as well as D2 and E2 are highlighted; G3 and H3 are not shown, as TNP computation failed for both of them (along with six other designs, 8/46 total).

We further retrospectively analyzed the experimentally validated designs across four properties: structure-prediction ranking, paratope composition, CDR3 conformation, and developability. To evaluate the ranking performance of structure prediction tools, we re-predicted all 46 V_H_H in complex with *α*-cobratoxin using ESMFold2 and compared ipTM and ipSAE min [30] to AF3 predictions (Fig. **2**C). Consistent with previous reports [12, 31], AF3 ipSAE min ranked the confirmed binders more favorably than AF3 ipTM (mean rank of D2/E2/G3/H3: 5.75 vs. 12.0 of 46). D2 and E2 ranked within the top 4 of 46 designs under both AF3 and ESMFold2 ipSAE min, whereas G3 and H3 ranked lower for AF3 ipSAE min (ranks 8–9 of 46) and even lower for ESMFold2 ipSAE min (ranks 16–23). Paratope composition analysis of AF3-predicted complexes suggested that interactions were primarily mediated by CDR3 residues, with smaller contributions from CDR1 and CDR2, consistent with typical V_H_H binding modes. The C-terminal 5 residues of *α*-cobratoxin were excluded from this analysis, as they were unresolved in reference structures and showed low predicted confidence in AF3 predictions. Notably, we observed a seemingly substantial contribution from framework residues (Fig. **2**D), while all four experimentally validated binders were still predicted to be predominantly driven by CDR3 binding. Consistent with the model comparison results, most selected designs adopted an extended CDR3 conformation. Using the *α*101 and *τ* 101-angle classification, 38/46 were classified as extended, 4/46 as kinked, and 4/46 as other. All four experimentally validated binders (D2, E2, G3, H3) were predicted to have an extended CDR3 conformation (Fig. **2**E). Finally, we assessed potential developability risks using the Therapeutic Nanobody Profiler (TNP), which flags V_H_Hs (green/amber/red) by comparison to properties of clinical-stage V_H_Hs. While most designs fell within the clinical-derived ranges across metrics, a subset were flagged, most frequently for charge-related properties (Fig. **2**F). Specifically, 8/38 of designs were classified as red for patches of positive charge (PPC) and 10/38 as red for patches of negative charge (PNC). The elevated PNC flags likely reflect the positively charged hotspot residues (R33, K35, R36) that define the target epitope. D2 and E2, like the parental framework 9GCN, pass all TNP metrics except PPC. The PPC flags in D2 and E2 were attributed to positively charged CDR residues outside the AF3-predicted paratope, suggesting that if optimization is required, these positions may be modifiable without affecting binding. We note that G3 and H3, alongside six other designs, failed the TNP assessment.

### Lead hits show favorable biophysical and developability profiles and present a distinct paratope

Since D2 and E2 showed the highest initial BLI response (Fig. S9), we considered these as our lead candidates and produced them at large-scale using *E. coli* SHuffle T7 expression. After C-tag purification, the two V_H_Hs were further purified by size exclusion chromatography (SEC), which indicated that D2 and E2 are predominantly monomeric, with monomer fractions of approximately 80.7% and 91.8%, respectively (Fig. **3**A). SDS-PAGE of the collected monomer fractions confirmed the expected molecular weight and high purity (Fig. S10).

**Figure 3.**
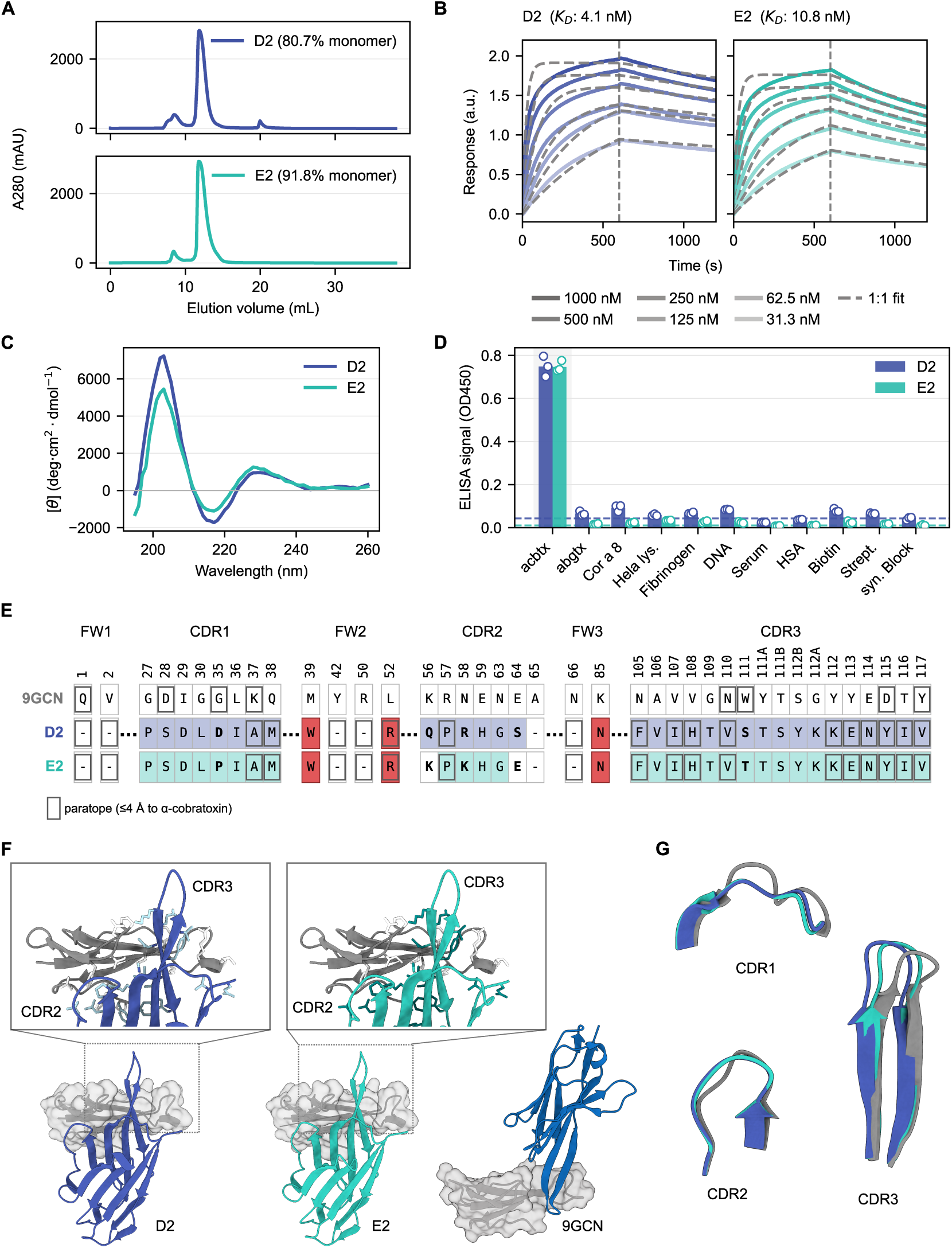
Biophysical, developability, sequence, and structural evaluation of lead V_H_Hs. **A)** Size-exclusion chromatography traces for the lead V_H_Hs D2 and E2. The main peak corresponds to the monomeric V_H_H fraction. **B)** Bio-layer interferometry analysis of D2 and E2 measured across six concentrations. Dissociation constants (*K_D_*) were estimated using a 1:1 binding model; fitted curves are shown as dashed lines. **C)** CD spectra confirming that D2 and E2 are *β*-sheet rich, consistent with the expected V_H_H secondary structure. **D)** Polyreactivity screening performed at 10 nM V_H_H concentration against a panel of nine off-targets, with *α*-cobratoxin included as a positive binding control. Binding was measured by ELISA, with wells coated with off-targets and bound V_H_Hs detected using an anti-Flag antibody. Dashed lines indicate background signal to a non-protein/non-DNA synthetic blocker control for D2 and E2. **E)** Sequence alignment of D2, E2, and the parental 9GCN framework. Dashes indicate amino acid identity with 9GCN; colored residues indicate positions that differ from 9GCN (red: framework mutations; other colors: CDR differences). Bold residues highlight differences between D2 and E2. Residues with grey outline indicate paratope residues. **F)** AF3-predicted structures of D2 and E2 in complex with *α*-cobratoxin, aligned on the toxin and compared to the parental 9GCN–*α*-cobratoxin complex. D2 and E2 are predicted to adopt a substantially different binding mode compared to 9GCN, approaching the epitope from a distinct angle and engaging a novel paratope surface. Zoomed insets shows that D2 and E2 share nearly identical predicted binding modes and interacting residues. **G)** Structural overlay of D2 and E2 CDR regions aligned to the corresponding 9GCN CDRs in grey.

Binding affinities were determined by measuring multiple V_H_H concentrations by BLI, which revealed that both V_H_Hs bind with *K_D_* values in the low nanomolar range: 4.1 nM (D2) and 10.8 nM (E2) (Fig. **3**B and Table S 3). This difference was driven almost entirely by the dissociation rate: D2 and E2 had comparable association rates (*k*_on_ = 4.28 *×* 10^4^ and 4.01 *×* 10^4^ M*^−^*^1^s*^−^*^1^, respectively), whereas the dissociation rate of E2 (*k*_off_ = 4.33 *×* 10*^−^*^4^ s*^−^*^1^) was *∼*2.5-fold faster than that of D2 (*k*_off_ = 1.75 *×* 10*^−^*^4^ s*^−^*^1^) (Table S 3). Circular dichroism (CD) confirmed that both V_H_Hs are *β*-sheet rich, showing the characteristic negative minimum near 217 nm and positive maximum near 203 nm expected for the immunoglobulin fold (Fig. **3**C). To obtain initial measurements for polyreactivity, we selected a panel of protein and non-protein targets and performed ELISA. At a V_H_H concentration of 10 nM, we observed minimal off-target binding, particularly for E2, with slightly elevated signals for D2 (Fig. **3**D). Notably, neither D2 nor E2 showed binding to *α*-bungarotoxin, a closely related long-chain *α*-neurotoxin that is also bound by the parental 9GCN V_H_H [24]. Despite D2 and E2 being flagged by TNP for patches of positive charge, neither showed strong binding to the negatively charged targets (HSA, dsDNA, fibrinogen). We note that at 100 nM V_H_H, the relative polyreactivity signal became higher but also showed elevated background signal (Fig. S11).

Sequence comparison to the original framework 9GCN showed minimal identity: D2 retains only one identical position in CDR2, while E2 shares three identical CDR2 positions with 9GCN; all other CDR residues differ. Notably, both D2 and E2 contain three framework mutations relative to 9GCN (Fig. **3**E). D2 and E2 stem from the same Germinal trajectory and represent different AbMPNN variants, differing at one position in CDR1, three in CDR2, and one in CDR3 (Fig. **3**E).

AF3 predictions of the V_H_H–*α*-cobratoxin complexes revealed that while D2 and E2 were predicted to bind to an overlapping epitope with 9GCN, their predicted binding mode is entirely different, approaching from the opposite side and presenting a novel paratope solution (Fig. **3**F). D2 and E2 display an almost identical predicted binding mode to each other since the residues predicted to interact with *α*-cobratoxin are identical (Fig. **3**E,F). Both designs also share the same number of interacting residues (19) and distribution across CDRs and framework residues (2/19 CDR1, 1/19 CDR2, 10/19 CDR3 and 6/19 framework). Notably, one of the introduced framework mutations (L52R) is predicted to be involved in binding. Despite sequence and paratope divergence from 9GCN, both designs converge on a similar CDR loop fold. After superposition on the framework backbone, the CDR2 C*α* RMSD to the 9GCN reference structure was 1.16 Å (D2) and 1.13 Å (E2), and the CDR3 RMSD was 1.96 Å (D2) and 1.34 Å (E2), both substantially lower than the corresponding CDR1 RMSD (2.84–2.86 Å) (Fig. **3**G). This close structural agreement for CDR3 is notable given its complete lack of sequence identity to 9GCN. This potentially indicates that the 9GCN framework can support this particular CDR3 loop geometry largely independent of its exact sequence.

### De novo designed V_H_Hs neutralize *α*-cobratoxin *in vitro* and *in vivo*

To assess the two lead candidates for functional *in vitro* neutralization, we performed patch-clamp experiments to record nAChR currents using a human-derived rhabdomyosarcoma cell line expressing muscle-type nAChRs. In the presence of a fixed concentration of *α*-cobratoxin (1.8 nM), we observed an IC_50_ of 4.2nM for D2 and 12.9nM for E2 (Fig. **4**A), meaning that E2 required a substantially larger molar excess of V_H_H over toxin to approach complete neutralization (*∼*188-fold vs. *∼*11-fold for 90% neutralization). Thus, D2 achieved more complete neutralization at substantially lower V_H_H-to-toxin ratios.

**Figure 4.**
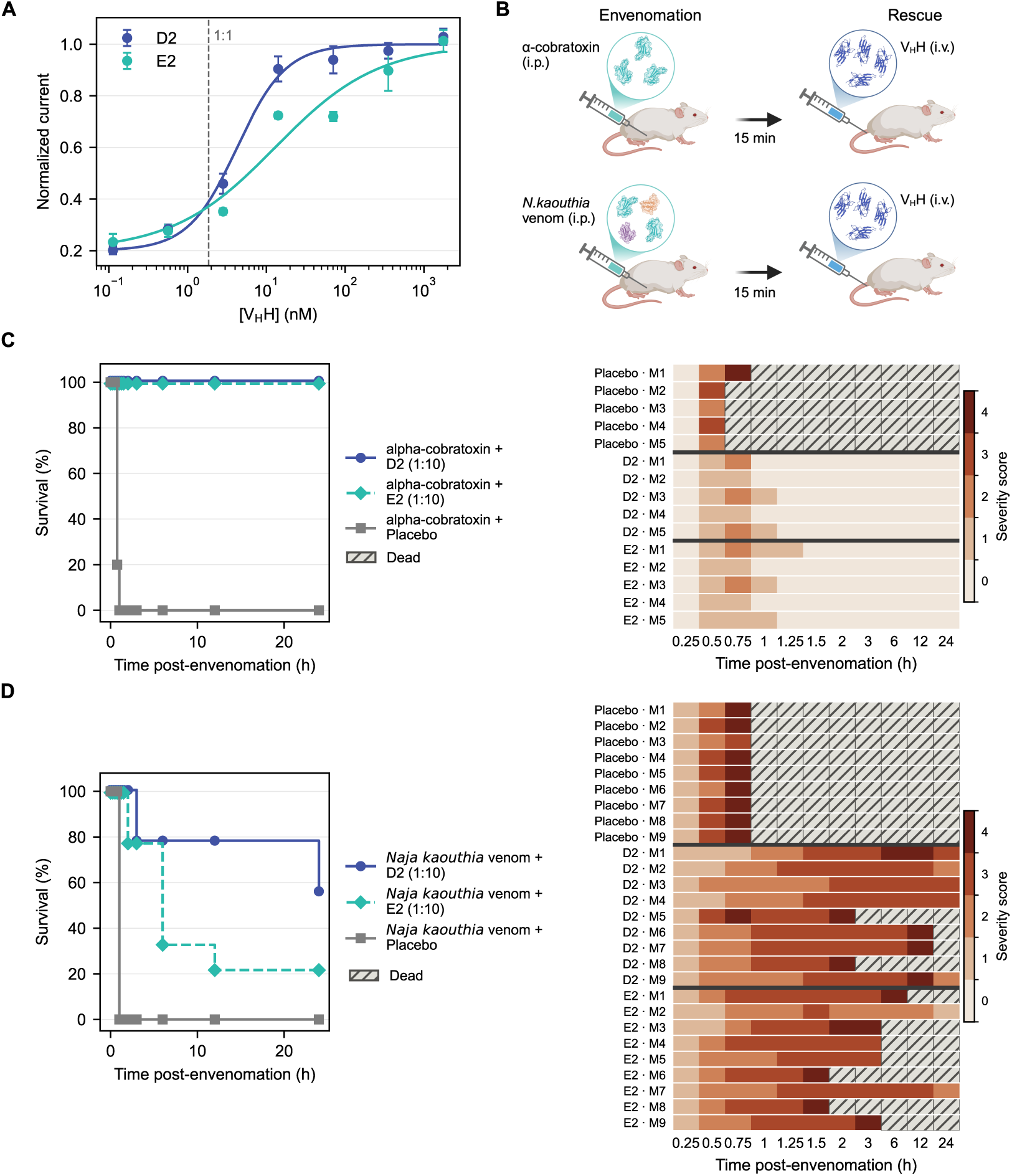
Functional *in vitro* and *in vivo* neutralization of. *α***-cobratoxin and whole *N. kaouthia* venom. A)** Patch-clamp dose-response curves for D2- and E2-mediated neutralization of *α*-cobratoxin-induced inhibition of muscle-type nAChR currents with the 1:1 molar V_H_H:toxin ratio indicated as a vertical dashed line. Data are shown as mean *±* SEM (*n* = 2–4 replicates per concentration). **B)** Overview of experimental set-up: mice (*n* = 5 for the *α*-cobratoxin challenge, *n* = 9 for the whole-venom challenge) were either challenged with *α*-cobratoxin or *N. kaouthia* venom by intraperitoneal (i.p.) injection. Rescue was performed 15 min later by injecting either D2 or E2 V_H_H by intravenous (i.v.) injection. **C)** *In vivo* protection against purified *α*-cobratoxin challenge. Left: survival curves for animals receiving toxin followed by D2, E2, or placebo (PBS). Right: clinical severity scores (0–4 scale) over the 24-h observation period for the same cohorts. *n* = 5 mice per cohort; V_H_Hs administered at 10-fold molar excess over toxin 15 min after challenge. **D)** As in **C)**, for animals challenged with whole *N. kaouthia* venom and *n* = 9 mice per cohort. M1 to M9 represent individual mice within each cohort.

Encouraged by these results, we evaluated the *in vivo* protection of D2 and E2 using a post-exposure rescue model (Fig. **4**B). Mice were challenged intraperitoneally with 3*×* LD_50_ of either purified *α*-cobratoxin or whole *N. kaouthia* venom, followed 15 min later by intravenous administration of D2 or E2 at a 10-fold molar excess over the toxin target (*n* = 5 per cohort). For the whole-venom challenge, the molar excess was estimated based on the reported *α*-cobratoxin content of *N. kaouthia* venom [22]. We included two control groups in the trial: To assess baseline lethality, placebo-treated mice received toxin or venom followed by PBS. To assess acute tolerability, a V_H_H-only control group received V_H_H in the absence of toxin or venom challenge. Clinical severity was scored on a 0–4 scale over 24 h (see Methods for scoring definitions).

Mice receiving V_H_H alone remained at baseline severity throughout the observation period, indicating acute tolerability of both candidates under test conditions (Fig. S12). Following challenge with purified *α*-cobratoxin, all placebo-treated animals succumbed within 45–60 min. In striking contrast, delayed administration of either D2 or E2 resulted in complete protection (5*/*5, 100% survival for both cohorts; Fisher’s exact test versus placebo, *p* = 0.0079 for each; log-rank test, *p* = 0.0016 for each). Treated animals developed only mild and short-lasting symptoms, reaching a peak mean severity of approximately 1.5*/*4 at 30–45 min before returning to baseline within 90 min (Fig. **4**C).

We next extended this to a rescue trial with whole *N. kaouthia* venom. Although *α*-cobratoxin is a major driver of the venom’s lethal neurotoxicity, whole venom contains additional neurotoxic and tissue-damaging components not targeted by the designed V_H_Hs, thereby representing a substantially more complex challenge. In this setting, all placebo-treated animals again succumbed rapidly, within 60 min. Both V_H_Hs significantly prolonged survival compared with placebo (log-rank, *p* = 3.7*×*10*^−^*^5^ for both). D2 additionally produced a significant improvement in 24-h survival compared with placebo (5/9 survivors; Fisher’s exact test, *p* = 0.029), whereas the corresponding improvement for E2 (2/9 survivors) was not significant (*p* = 0.47). Although D2 showed numerically greater protection than E2, the direct difference between the two candidates did not reach statistical significance (log-rank, *p* = 0.11). Clinical severity was greater and more persistent than following purified *α*-cobratoxin challenge, and surviving animals did not fully return to baseline within the 24-h observation period. The stronger protection observed with D2 is consistent with its higher affinity, slower off-rate, and steeper *in vitro* neutralization profile. Together, D2’s protection against both purified *α*-cobratoxin and whole venom identifies this de novo-designed V_H_H as a promising therapeutic lead.

## Discussion

Single-domain antibodies (V_H_Hs) offer a clinically precedented, antibody-based alternative to new-to-nature de novo scaffolds, but it has been unclear whether current de novo methods can reliably generate functionally relevant V_H_H binders. Here, using *α*-cobratoxin as a medically relevant target, we demonstrate that de novo V_H_H design pipelines can produce high-affinity binders with favorable therapeutic properties and *in vivo* protection. Further, a head-to-head comparison of three design models under standardized conditions reveals substantial differences between methods and frameworks; indicating that model choice and workflow design remain critical determinants of experimental success.

In our *in silico* comparison of Germinal [12], RFantibody [11], and BoltzGen [13], the diffusion-based methods achieved higher raw throughput, but Germinal (which relies on more computationally intensive hallucination) produced a substantially larger fraction of designs passing combined ipTM and RMSD self-consistency filtering (based on both AF3 and ESMFold2 re-prediction), and consequently a lower compute time per successful design. Self-consistency was a crucial complementary criterion, since many designs achieved reasonable ipTM scores but failed to maintain their binding mode upon re-prediction, which is consistent with recent reports [11].

Beyond confidence metrics, we observed method-dependent differences in CDR3 sequence novelty and loop conformation, indicating that current generative antibody design pipelines sample distinct regions of sequence and structural space even when key design inputs are held constant. Framework choice also influenced experimental outcomes: binders were obtained only from scaffolds with extended CDR3 conformations, while the 3EAK scaffold, used in both the Germinal and RFantibody studies, did not yield binders, though only eight V_H_Hs from this framework were tested. This underscores the potential value of evaluating multiple frameworks in prospective V_H_H design campaigns. Supporting the importance of CDR3 conformation, the lead candidates D2 and E2 adopted a CDR3 backbone closely matching the parental 9GCN V_H_H despite sharing no CDR3 sequence identity.

From a development perspective, most selected designs expressed as soluble protein (42/46), and we identified four binders, including two low-nanomolar leads (D2, E2), demonstrating that de novo V_H_H design can reach affinity regimes relevant for functional neutralization. Both leads showed favorable developability by SEC and CD, with high monomer content (80.7% and 91.8% for D2 and E2, respectively) and the expected *β*-sheet-rich secondary structure. While TNP flagged both leads for patches of positive surface charge (PPC), neither showed clear binding to negatively charged off-targets, suggesting these flags may not translate into functional polyreactivity. The modestly lower monomer fraction and slight polyreactivity signal for D2 indicate areas for further characterization.

Importantly, both leads represent genuinely novel binding solutions despite being derived from the parental 9GCN framework. Although the 9GCN V_H_H is itself known to bind *α*-cobratoxin, D2 and E2 share essentially no CDR sequence identity with it, and their AF3-predicted complexes suggest a substantially different binding mode, approaching the toxin from the opposite side and engaging a distinct paratope. Further supporting distinct recognition, neither D2 nor E2 bound to *α*-bungarotoxin, despite the parental 9GCN V_H_H being cross-reactive with this toxin [24]. Experimental structures would be needed to resolve this fully, though AF3 predictions of designed antibody complexes are generally considered close to experimental structures once a design is confirmed to bind [11, 12]. Notably, the two additional lower-affinity binders (G3 and H3) were obtained from the unrelated 7XL0 framework, showing that successful binding was not restricted to the parental *α*-cobratoxin-binding scaffold.

The central result of this study was that these de novo-generated antibodies not only bound *α*-cobratoxin with high affinity but neutralized it functionally, both *in vitro* and *in vivo*, achieving complete protection against purified toxin and partial protection against whole *N. kaouthia* venom. Notably, D2 and E2 differ 2.6-fold in affinity, attributable to a *∼*2.5-fold slower off-rate of D2. This was reflected in the patch-clamp data, where D2 required a lower molar excess for complete neutralization, and potentially in the whole-venom challenge, where D2 protected more animals (56% vs. 22%). This numerical divergence emerged against whole venom but not purified toxin, potentially because venom dosing relied on an estimated *α*-cobratoxin content; given the substantial variability in *N. kaouthia* venom composition [22], the true molar excess may have been well below the nominal 10-fold. We note, however, that the difference between D2 and E2 is not statistically significant, and the possibility that the two leads are functionally equivalent against whole venom cannot be excluded. Our results parallel the recent demonstration that de novo non-antibody binders neutralize *α*-cobratoxin [7], with comparable affinities, *in vivo* protection, and tolerability. We extend this to whole *N. kaouthia* venom, where the difference between D2 and E2 underscores the importance of whole-venom testing. Whether antibody-based or non-antibody-based formats will ultimately prove advantageous for antivenoms or protein therapeutics broadly remains open and may be case-dependent. The current value of the V_H_H approach lies in leveraging a clinically validated modality with established routes to optimization and approval.

Several limitations remain. Our model and framework comparisons were performed against a single target and across only three scaffolds, and because RFantibody and BoltzGen did not yield designs passing our filtering criteria even at increased sampling, the comparison between methods remained entirely *in silico*. This limits how far we can generalize our conclusions about relative model performance, though it does not diminish the value of the comparison framework itself as a template for evaluating design tools under shared, prospective conditions. While our results are consistent with the idea that arbitrary shuffling of CDRs between frameworks is not generally well tolerated [26], testing additional frameworks against additional targets will be needed to draw firm conclusions. A further caveat is that 9GCN is neither humanized nor otherwise optimized for developability, unlike frameworks commonly used in de novo V_H_H design; although both leads were well tolerated acutely *in vivo* and immunogenicity assessment would ultimately be required regardless of framework choice, since CDR sequences themselves can be a major determinant of immunogenicity [32]. We also note that 6 of 19 predicted paratope residues for both leads are framework residues; while elevated framework involvement in the paratope is a common feature of V_H_Hs [33], it may complicate downstream humanization, as framework positions contributing to binding cannot be freely modified without risking loss of affinity [34].

We therefore view this workflow as a route to a strong therapeutic starting point rather than a finished molecule. Notably, D2 and E2 differ by only a small number of AbMPNN-refined positions, yet show clear differences in affinity and neutralization, indicating that an initial design hit could be further improved using the same design framework. Taken together, our results suggest that generative binder design can now contribute meaningfully to therapeutic antibody discovery, rather than being limited to non-antibody scaffolds with little clinical precedent. More broadly, by integrating recent methodological advances in an end-to-end pipeline, de novo V_H_H design can support the rapid generation of candidates for applications such as recombinant antivenom development.

## Materials and Methods

### Model comparison setup and evaluation

Model comparisons were performed at the stage in each pipeline immediately preceding its structure-prediction-based scoring step, i.e. the point at which designs would normally get re-predicted and ranked with a structure prediction model. For Germinal, we used sequences produced by each generative trajectory prior to AbMPNN redesign. Importantly, we retained sequences from all trajectories regardless of their success at intermediate filtering steps (logits, softmax, and semi-greedy phases), to avoid bias from Germinal’s built-in quality filters. For RFantibody and BoltzGen, we used the sequences output by their default inverse-folding steps (ProteinMPNN and BoltzIF, respectively) together with the corresponding designed structures. Three V_H_H scaffolds were used as input across all models: TPL1158 01 C09 (PDB ID: 9GCN), NbBCII10 humanized (PDB ID: 3EAK), and Vobarilizumab (PDB ID: 7XL0). CDR boundaries and lengths were identified with AbNumber using IMGT numbering and provided to each model such that CDR lengths were fixed to match the parental scaffold. A full-length AlphaFold3 (AF3)-predicted *α*-cobratoxin structure was used as the common target input for all models. All methods were provided the same target hotspot residues (positions D27, R33, K35, R36, and V37), selected based on key epitope residues reported previously [24].

All designed candidates were re-predicted as V_H_H–toxin complexes using AF3. AF3 was run with a single seed and five diffusion models per seed, and ipTM was taken from the top-ranked model. Multiple sequence alignments (MSAs) were only provided for the target sequence and generated with ColabFold MMseqs2 [35]. To assess structural plausibility, we applied a self-consistency filter based on the root-mean-square deviation (RMSD) between the designed V_H_H and its AF3 re-predicted V_H_H structure after superposition of the complexes on the target; RMSD was computed over V_H_H C*α* atoms.

For cross-validation, ESMFold2 predictions were generated without MSA for the V_H_H sequence and with precomputed target MSA (same as AF3 setup), using a single seed and default parameters (20 loops, 100 sampling steps). ipTM scores were extracted from the top-ranked model.

Design speed comparison was performed by running each method once per scaffold against *α*-cobratoxin on an NVIDIA L40S GPU. For Germinal, 100 designs were generated per scaffold, whereas 300 designs per scaffold were generated for RFantibody and BoltzGen. We report two complementary runtime comparisons. First, raw throughput, defined as the number of designs generated per unit wall-clock time, computed either for the design stage alone or including AF3 re-prediction time for all designs (design+AF3). Second, time per successful design, defined as the design-stage wall-clock time divided by the number of designs meeting a specified success criterion (an ipTM threshold, with or without the RMSD self-consistency filter).

Software versions used for the computational analyses were AlphaFold3 v3.0.1 (AlphaFold3), BoltzGen v0.2.0 (BoltzGen), and ESMFold2 v3.4.0 (ESMFold2). Germinal was run at commit 88d7f85 (Germinal), and RFantibody at commit 8d9d402 (RFantibody).

### CDR3 novelty profiling

For each designed V_H_H, we quantified CDR3 novelty by aligning the full-length V_H_H sequence against the Structural Antibody Database (SAbDab) [36] using MMseqs2 [37]. For each design, we identified the database hit with the highest CDR3 sequence identity and reported that identity as the novelty metric. We performed an analogous analysis against the Observed Antibody Space (OAS) [38] and INDI [39] V_H_H-only sequence databases, requiring a minimum sequence identity of 60% and minimum coverage of 90% to keep the search computationally tractable against these much larger databases.

### Sequence-space embedding by multidimensional scaling (MDS)

To visualize how designs distribute in sequence space, we embedded all generated CDR3 sequences together with naturally occurring CDR3s from an immunized V_H_H library screened against *α*-cobratoxin [27] using metric multidimensional scaling (MDS). Pairwise distances were computed as Levenshtein (edit) distances between CDR3 amino-acid strings. The resulting all-by-all distance matrix was projected into two dimensions with metric MDS (precomputed dissimilarities; sklearn.manifold.MDS), and the 2D coordinates were used for plotting, where proximity reflects higher sequence similarity under the edit-distance metric.

### CDR3 conformation profiling

CDR3 loop conformation was classified using a geometric scheme based on the *α*_101_ dihedral angle and *τ*_101_ bond angle computed from C*α* atoms at Chothia positions 100x–103 [28]. Designs were categorized as kinked, extended, or other according to the published threshold ranges: kinked was defined as 0*^◦^ < α*_101_ *<* 120*^◦^* and 85*^◦^ < τ*_101_ *<* 130*^◦^*; extended was defined as *α*_101_ *< −*100*^◦^*and 100*^◦^ < τ*_101_ *<* 145*^◦^*; all remaining designs were classified as other. Because *α*_101_ is a dihedral angle reported on a (*−*180*^◦^,* 180*^◦^*] branch cut, the extended criterion was additionally satisfied for *α*_101_ *>* 150*^◦^* to capture designs whose dihedral values wrap around this seam without representing a distinct fold.

### Large-scale de novo V_H_H design using Germinal

Crystal structure for *α*-cobratoxin was derived from 9GCN and served as the target input in this V_H_H design campaign. The same epitope hotspot residues used for the model comparison were used. Designs generated against *α*-cobratoxin for each scaffold were filtered using the default Germinal V_H_H design criteria including RMSD *<* 6.0 Å, ipTM *>* 0.80, and pTM *>* 0.80 (see all initial and final filters in Table S2). The only modification to default Germinal run settings was a change in the minimum number of hotspot residues required to be targeted by the CDRs, which was reduced from three to two. All designs passing this filtering step, along with AbMPNN-redesigned V_H_H structures, were subsequently re-predicted using AF3 and the full-length *α*-cobratoxin amino acid sequence, with MSA generation performed for both the V_H_H and the target. Final design filtering was based on RMSD *<* 6.0 Å, ipTM *>* 0.80, and pTM *>* 0.80. A total of 48 designs were selected for gene synthesis, including 8 designs from the 3EAK scaffold and 20 designs each from the 9GCN and 7XL0 scaffolds. These designs were profiled for CDR novelty and CDR3 conformation as described above. Epitope and paratope residues were defined as residues containing at least one heavy atom within 4 Å of any heavy atom on the interaction partner. Developability metrics were obtained using the online Therapeutic Nanobody Profiler (TNP) [40]; a subset of submissions failed to return results and are reported as missing values.

### Golden Gate cloning and transformation into SHuffle T7 cells

Designed protein sequences were reverse-translated and codon-optimized using the GenScript GenSmart Codon Optimization tool for expression in *Escherichia coli*. Codon-optimized gene sequences were generated with flanking sequences compatible with Golden Gate cloning and PCR amplification. V_H_H gene fragments were cloned into a custom pET expression vector containing BsaI sites, a C-terminal FLAG tag, and a C-tag by Golden Gate assembly with BsaI-HFv2 (New England Biolabs) and T4 DNA ligase (Thermo Fisher). Reactions (10 µL) contained 75 ng vector and 35 ng insert, 1 µL 10*×* T4 ligase buffer, 1 µL T4 DNA ligase, and 1 µL BsaI-HFv2, with water to volume. Cycling was performed as 45 cycles of 37 °C (5 min) and 16 °C (5 min), followed by 37 °C (5 min) and heat inactivation at 80 °C (10 min). For transformation, 5 µL of the Golden Gate reaction was added to 50 µL chemically competent SHuffle T7 cells (New England Biolabs), incubated on ice for 30 min, heat shocked at 42 °C for 30 s, and returned to ice for 5 min. SOC medium (150 µL) was added and cells were recovered at 30 °C for 1 h before plating on selective agar and incubation at 30 °C for 36 h.

### Small-scale protein expression

Single colonies were picked from selection plates into 1 mL 2*×*YT medium supplemented with kanamycin in 96-deep-well plates and grown overnight at 30 °C with shaking (800 rpm, 80% humidity). The next day, 4 *×* 10 µL of each overnight culture was used to inoculate 4 mL fresh 2*×*YT + kanamycin (1 mL in each of four 96-deep-well plates). Cultures were grown to OD_600_ ≈ 0.5 and induced with IPTG (final 1 mM). Induced cultures were incubated overnight at 25 °C with shaking (800 rpm).

### C-tag purification and protein quantification

Induced cultures in 96-well plates were pelleted (4000*×g*, 10 min) and the supernatant was removed. Pellets were frozen at *−*20 °C for 30 min, resuspended in 400 µL lysis buffer per well, incubated for 30 min at room temperature with shaking, and clarified by centrifugation (7000*×g*, 20 min, 4 °C). CaptureSelect^TM^ C-tagXL affinity resin (Thermo Fisher) was equilibrated on a vacuum manifold with three washes of 700 µL TBS (50 mM Tris, 150 mM NaCl, pH 7.4). Clarified lysate (400 µL) was loaded per column and incubated for 30 min with shaking. Columns were washed three times with 700 µL TBS, and bound V_H_Hs were eluted twice with 180 µL glycine (100 mM, pH 2.7; 5 min incubation each). Eluates were collected into 20 µL neutralization buffer (1 M Tris, 1.5 M NaCl, pH 8.5) by centrifugation (2000*×g*). Columns were regenerated with three washes of 700 µL elution buffer, incubated with 6 M urea for 5 min, washed three times with TBS, and stored in 20% ethanol. Protein concentrations were quantified using the Qubit^TM^ Protein Assay (Thermo Fisher). Working solution was prepared by mixing reagent and buffer 1:200. Standards (10 µL + 90 µL working solution) were measured in triplicate (25 µL per well), and samples were measured by mixing 2.5 µL eluate with 22.5 µL working solution in a black 384-well plate. Fluorescence was read on a Victor Nivo plate reader after 3 min shaking.

### Flow-induced dispersion analysis (FIDA)

*α*-cobratoxin was fluorescently labeled with Alexa Fluor^TM^ 647 (Thermo Fisher) according to the manufacturer’s protocol. Excess label was removed using Dye Removal Columns (Thermo Fisher). For screening, C-tag–purified V_H_Hs were pre-mixed with Alexa Fluor 647–labeled *α*-cobratoxin and diluted in PBS + 0.03% Tween-20 to final concentrations of 50 nM V_H_H and 125 nM toxin. Because the screen was qualitative and based on the change in *R_h_* of the labeled toxin, sub-stoichiometric V_H_H was sufficient to detect complex formation. Binding measurements were performed on a FIDA 1 instrument (Fida Biosystems ApS) using LED-induced fluorescence detection (excitation 640 nm) and permanently coated capillaries to reduce adsorption. The premix format was used throughout, and measurements were acquired as single runs. Labeled *α*-cobratoxin without V_H_H was included as a control and to estimate the hydrodynamic radius (*R_h_*) of the target. Candidates were classified as screening hits when addition of V_H_H produced a clear increase in the apparent hydrodynamic radius (*R_h_*) of labeled *α*-cobratoxin relative to the toxin-only control, consistent with complex formation. Each measurement consisted of an automated sequence: (1) 1 M NaCl wash (30 s, 3500 mbar), (2) H_2_O wash (30 s, 3500 mbar), (3) equilibration in PBS with 0.03% Tween-20 (20 s, 2000 mbar), (4) injection of PBS with 0.03% Tween-20 (30 s, 2000 mbar), (5) injection of the pre-mixed V_H_H–toxin sample (10 s, 50 mbar), and (6) injection of PBS with 0.03% Tween-20 (200 s, 400 mbar). Results were analyzed with custom Python scripts.

### Large-scale expression and SEC

The plasmids of D2 and E2 were confirmed by sequencing (Eurofins Genomics). 2*×*YT cultures were inoculated with the respective clone, induced with IPTG, and incubated overnight at 20 °C. The culture was spun down, the pellet resuspended in TBS (pH 7.4) containing 0.05 µL/mL benzonase, 1 mM MgCl_2_ and cOmplete EDTA-free protease inhibitors (Roche), and lysed using sonication. The lysate was spun down, the supernatant filtered through a 0.45 µm syringe filter, and purified on an NGC BioRad system using a CaptureSelect^TM^ C-tagXL prepacked 5 mL column (Thermo Fisher Scientific). TBS served as wash buffer and 2 M MgCl_2_ as elution buffer. The column was stripped with 0.1 M glycine (pH 2). Protein was concentrated on a 10 kDa MWCO Amicon centrifugal filter (Sigma Aldrich) and applied to a Superdex 75 10/300GL size exclusion chromatography (SEC) column (Cytiva) in TBS on an NGC BioRad system. The size of both V_H_Hs was confirmed by SDS-PAGE.

### Bio-layer interferometry (BLI)

Octet^®^ streptavidin (SA) biosensors were equilibrated in HEPES buffer (10 mM HEPES, 150 mM NaCl, 3 mM EDTA, 50 mM MES, 0.05% Tween-20, pH 7.2) for 30 min prior to analysis. Biotinylated *α*-cobratoxin was diluted in the same buffer to 200 nM. Measurements were conducted on an Octet RED96e system (ForteBio) at 30 °C. Biotinylated *α*-cobratoxin was immobilized onto SA biosensors to a loading response of *∼*1.0 nm. Association was performed for 600 s followed by dissociation for 600 s in HEPES buffer. Data were reference-subtracted using an antigen-loaded SA biosensor dipped into buffer in the absence of V_H_H. One-point measurements of D2, E2, G3, and H3 were performed at 200 nM V_H_H concentration. Large-scale produced D2 and E2 were titrated in six two-fold dilutions from 1000 nM to 31.3 nM. Curves were analyzed using Octet Analysis Studio v12.2.2.26 (ForteBio) and custom Python scripts. The curves were fitted with a 1:1 model.

### Polyreactivity screening

MaxiSorp 96-well plates were coated overnight at 4 *^◦^*C with 10 *µ*g/mL human serum albumin (HSA; #A1653, Sigma-Aldrich), biotin (#B4501, Sigma-Aldrich), streptavidin, bovine fibrinogen (#F8630, Sigma-Aldrich), recombinant Cor a 8, *α*-cobratoxin, *α*-bungarotoxin (in-house), or HeLa cell lysate (in-house). Double-stranded DNA (dsDNA; #Q33230, Qubit) was coated at 5 *µ*g/mL, and human serum (#H6914, Sigma-Aldrich) was diluted 1:50 prior to coating. Plates were washed three times with PBS containing 0.05% Tween-20 (PBST) and blocked with 300 *µ*L Protein-Free Blocking Buffer (#37570, Thermo Fisher Scientific) for 1 h at room temperature (RT). Following three washes with PBST, 50 *µ*L of V_H_H diluted to 10 nM or 100 nM in PBS containing 0.5% BSA was added to each well and incubated for 1 h at RT. Plates were washed three times with PBST before incubation with anti-FLAG-HRP diluted 1:10,000 in PBS containing 0.5% BSA for 1 h at RT. After a final three washes with PBST, 50 *µ*L of TMB substrate (#34021, Thermo Fisher Scientific) was added and incubated until color development. The reaction was stopped by addition of 50 *µ*L of 0.5 M H_2_SO_4_, and absorbance was measured at 450 nm using a VICTOR Nivo plate reader (PerkinElmer).

### Circular dichroism

The secondary structure of recombinant V_H_Hs was analyzed by far-UV circular dichroism (CD) spectroscopy using a Jasco J-715 spectropolarimeter with a 0.1 cm pathlength quartz cuvette. Purified V_H_Hs were buffer-exchanged into PBS (pH 7.4) and spectra were recorded from 190 to 260 nm at 25 °C. Three consecutive scans were averaged, the corresponding PBS spectrum was subtracted, and the data are presented as mean residue ellipticity (deg*·*cm^2^*·*dmol*^−^*^1^).

### *In vitro* neutralization using electrophysiology

Human-derived rhabdomyosarcoma RD cells (American Type Culture Collection) endogenously expressing the muscle-type nAChR ((*α*_1_)_2_*β*_1_*γδ*) were used for electrophysiology experiments, as previously described [41]. Whole-cell planar patch-clamp was performed on a Qube automated electrophysiology platform (Sophion Bioscience). *α*-cobratoxin was used at a single fixed concentration of approximately 1 IC_80_ (1.85 nM), obtained by titration of the toxin. V_H_Hs D2 and E2 were added in a seven-point, five-fold dilution series (0.11–1765 nM) at this fixed toxin concentration; buffer-only wells served as a negative control. The ability of the toxin to inhibit an acetylcholine (ACh, 70 µM) response in the presence of binder was normalized to the full ACh response and averaged across *n* = 2 *−* 4 technical replicates per concentration–condition pair. Concentration-response curves were fit with a four-parameter logistic (Hill) model using a custom Python script to estimate IC_50_ values for each binder-toxin pair, with the bottom and top asymptotes fixed at the assay-defined values of 0.2 (residual normalized current with toxin alone, corresponding to *∼*1 IC_80_ inhibition) and 1.0 (full, uninhibited ACh response), respectively.

### *In vivo* experiments

Assays used male non-Swiss albino mice (20–24 g); all doses were adjusted to body weight.

Purified *α*-cobratoxin (7,820 Da) and whole *N. kaouthia* venom were obtained from Latoxan S.A.S. Toxin/venom was solubilized in PBS at 1.0 mg mL*^−^*^1^ and diluted in PBS as needed.

The LD_50_ of purified *α*-cobratoxin (i.p., mouse) was taken as 0.294 *µ*g g*^−^*^1^ [18]. The LD_50_ of whole *N. kaouthia* venom (i.p., mouse) was taken as 0.148 *µ*g g*^−^*^1^ [42].

Mice were challenged intraperitoneally with 3*×* LD_50_ of either purified *α*-cobratoxin or whole *N. kaouthia* venom, administered as a 100 *µ*L bolus in the right lower abdominal region. Fifteen minutes after challenge, animals received D2 or E2 intravenously via the left lateral caudal vein in a volume of 200 *µ*L, dosed at a 10-fold molar excess over toxin. For the purified *α*-cobratoxin challenge, this excess was calculated directly from the administered *α*-cobratoxin dose. For the whole-venom challenge, the *α*-cobratoxin content of the venom dose was estimated as 32.3% of total venom protein by mass [22], and the V_H_H dose was calculated as a 10-fold molar excess over this estimated *α*-cobratoxin amount. For the purified *α*-cobratoxin challenge, the following cohorts were run: (i) toxin followed by D2 (*n* = 5), (ii) toxin followed by E2 (*n* = 5), (iii) toxin followed by PBS (placebo, *n* = 5), and (iv) D2 alone and (v) E2 alone without toxin (*n* = 3 each), the latter two to assess acute tolerability. For the whole-venom challenge, the D2, E2 and placebo cohorts were extended to *n* = 9 by pooling two independent trials to enable a more robust comparison.

Clinical severity was scored on a 0–4 scale by an observer at 0.25, 0.5, 0.75, 1, 1.25, 1.5, 2, 3, 6, 12 and 24 h post-challenge (Table 1). The scale was adapted for murine models from an established equine grading scale, with a related published adaptation for envenomation available for porcine models [43].

**Table 1:** Clinical severity scoring criteria for *in vivo* challenge experiments.

| Score | Description |
| --- | --- |
| 0 | Mice alert and active as cage is opened for observation, and are responsive upon stimulation. |
| 1 | Mice aware but not active as cage is opened for observation; generally able to move around during passive observation, or responsive upon stimulation, but showing beginning signs of discomfort. |
| 2 | Mice readily respond to stimulation but are unwilling to move once stimulation has ceased; signs of discomfort such as excessive grooming and hunched posture are observable and obvious. |
| 3 | Mice inactive throughout passive observation, require multiple attempts at stimulation to move, movements are slow with abnormalities in gait and carriage. |
| 4 | Mice lying flat/recumbent as cage is opened for observation; multiple attempts at stimulation result in 0–4 steps before lying flat or on side. Any animal that can take more than 4 steps upon stimulation or during passive observation will not receive a score of 4. |

Protection was assessed as percent survival at 24 h, together with the clinical severity time course for each cohort.

### Statistical analysis

Survival outcomes were evaluated using both endpoint and time-to-event analyses. Differences in 24-h survival proportions between treatment and placebo groups were assessed using Fisher’s exact test. Time-to-death distributions were compared using the log-rank (Mantel–Cox) test. For the purified *α*-cobratoxin challenge, D2 and E2 treatment groups were each compared with placebo. For the whole-venom challenge, D2 and E2 were each compared with placebo, and the two treatment groups were also compared directly. The whole-venom cohorts (*n* = 9 per treatment group) were pooled from two independent trials. A two-sided *p <* 0.05 was considered statistically significant. Survival analyses were performed in Python using the lifelines package.

### Ethics statement

All mice utilized in this study were handled in accordance with protocol 2508D-SM-SMLBirds-28 approved by the University of Northern Colorado Institutional Animal Care and Use Committee (to SPM).

### Data Availability

All data supporting the findings of this study are provided as Supplementary Material, including a summary of the 48 designed V_H_Hs (sequences, expression, binding, developability, and biophysical characterization data) and AlphaFold3 structure predictions for all designs.

## Supporting information

VHH analysis and AF3 predictions

## Acknowledgments

We would like to thank the HPC support team at the DTU Computing Center (DCC) for their valuable assistance in setting up the required infrastructure for this work. We also thank Soumik Ray (DTU Bioengineering) and Kritika Ray (Fida Biosystems) for their help in establishing the FIDA assay. We further thank Pia Haugaard Nord-Larsen and the administration team at DTU Bioengineering for their continuous support.

## Funding

T.P.J. and M.D.O. acknowledge support from the Alliance programme under the EuroTech Universities agreement. M.B.V. acknowledges support from Independent Research Fund Denmark (grant ID 10.46540/4283-00192B). NIVI Research Center is part of The Novo Nordisk Foundation Initiative on Vaccines and Immunity (NIVI) and is supported by The Novo Nordisk Foundation (Grant number: NNF23SA0088562). D.R.E. acknowledges support from Novo Nordisk A/S as part of the DTU Novo Nordisk Closed-Loop Optimization initiative. S.P.M. and S.K. are supported by a grant from the National Science Foundation (IOS-2307044).

## Competing Interests

T.P.J. is a co-founder and shareholder of AffinityAI ApS. E.V.S.L. is an employee of AffinityAI ApS. The company had no role in the design, execution, interpretation, or funding of this study. All other authors declare they have no competing interests.

## Author contributions

M.D.O. and E.V.S.L. performed the model-comparison analyses. C.P.J. performed the multidimensional scaling analysis and assisted with CDR3 loop conformation assignment. E.V.S.L. led the large-scale Germinal design campaign. M.D.O., D.R.E., and E.V.S.L. performed the initial binding screening. K.H.B. performed the large-scale expression, SEC and BLI. M.B.M. conducted the CD and polyreactivity screening. K.H.B. K.B. and M.B. performed the patch-clamp experiments. M.F.Q. performed the structural characterization. S.K. and S.P.M. performed the *in vivo* experiments. T.P.J. supervised the project. M.D.O. and T.P.J. wrote the original draft of the manuscript. All authors reviewed and approved the final manuscript.

## Supplementary Figures

**Fig. S1.**
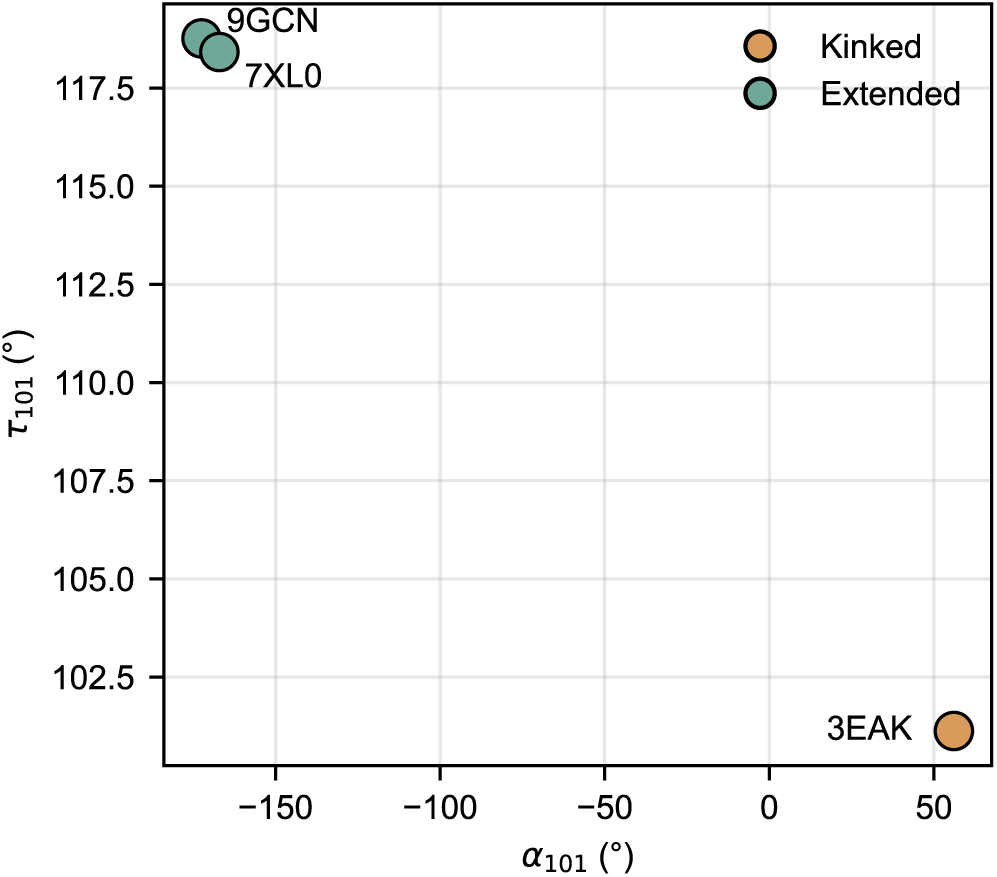
CDR3 conformation of selected V_H_H frameworks: *α*101 and *τ* 101 angles for the three scaffolds used for design (9GCN, 7XL0, 3EAK). 9GCN and 7XL0 adopt an extended CDR3 conformation, whereas 3EAK adopts a kinked conformation.

**Fig. S2.**
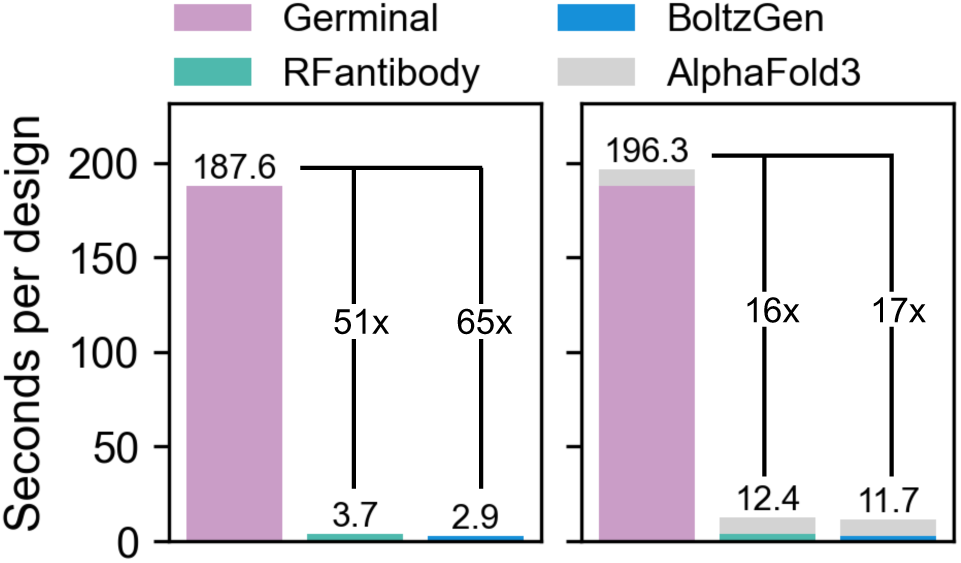
Runtime without success filtering: Seconds per design for Germinal, RFantibody, and BoltzGen during V_H_H generation, reported with and without AlphaFold3 (AF3) structure prediction. Fold-changes are annotated relative to Germinal.

**Fig. S3.**
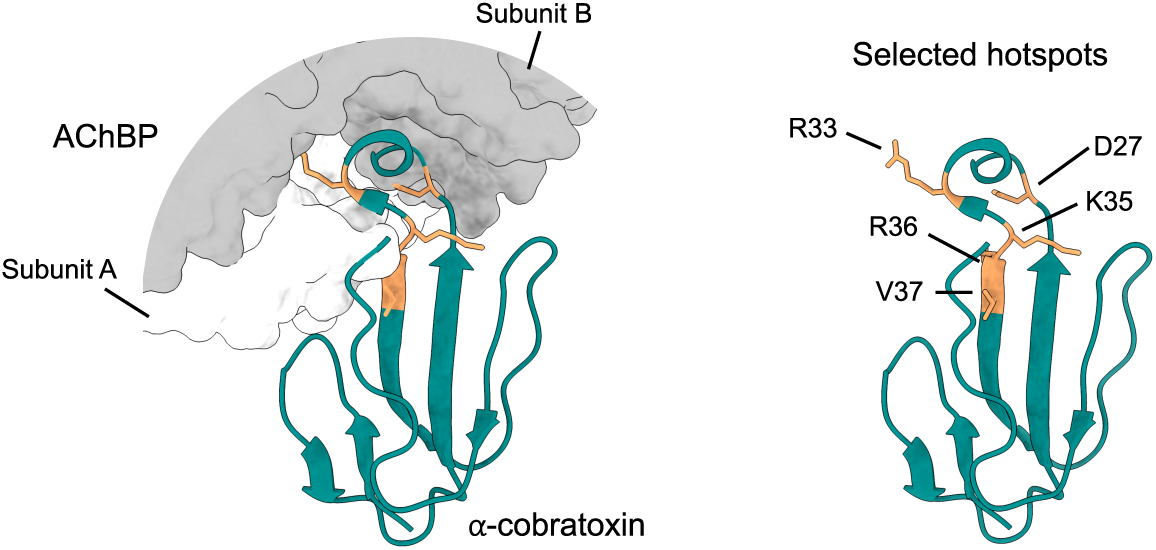
Hotspot definition: Structure of *α*-cobratoxin (teal) bound to acetylcholine-binding protein (AChBP; PDB ID 1YI5 [44]). Two AChBP subunits (grey and white) form the binding site for the toxin. Residues selected as design “hotspots” are highlighted in orange, all engaging in AChBP binding.

**Fig. S4.**
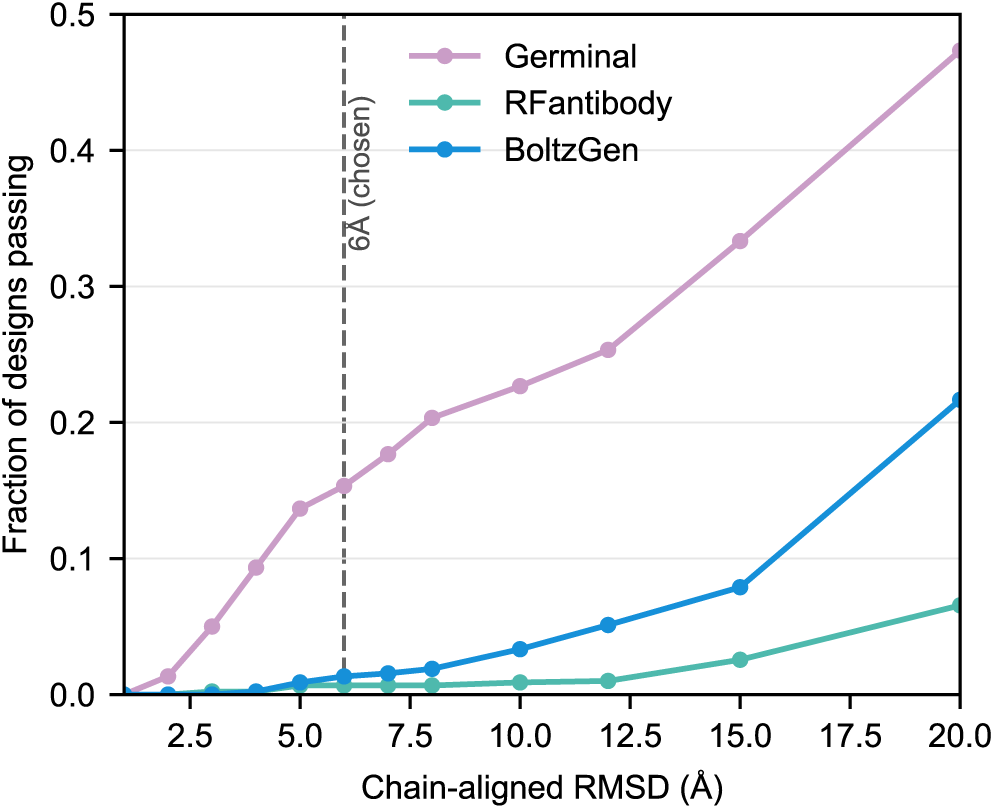
RMSD self-consistency threshold sensitivity: Fraction of designs per method passing the AF3 self-consistency filter (chain-aligned V_H_H RMSD after target-chain alignment) as the threshold is relaxed from 1 to 20 Å. At any threshold in the 2–5 Å range, RFantibody and BoltzGen have essentially zero passing designs; 6 Å (dashed line) is the loosest threshold that still yields non-trivial pass counts for all three methods.

**Fig. S5.**
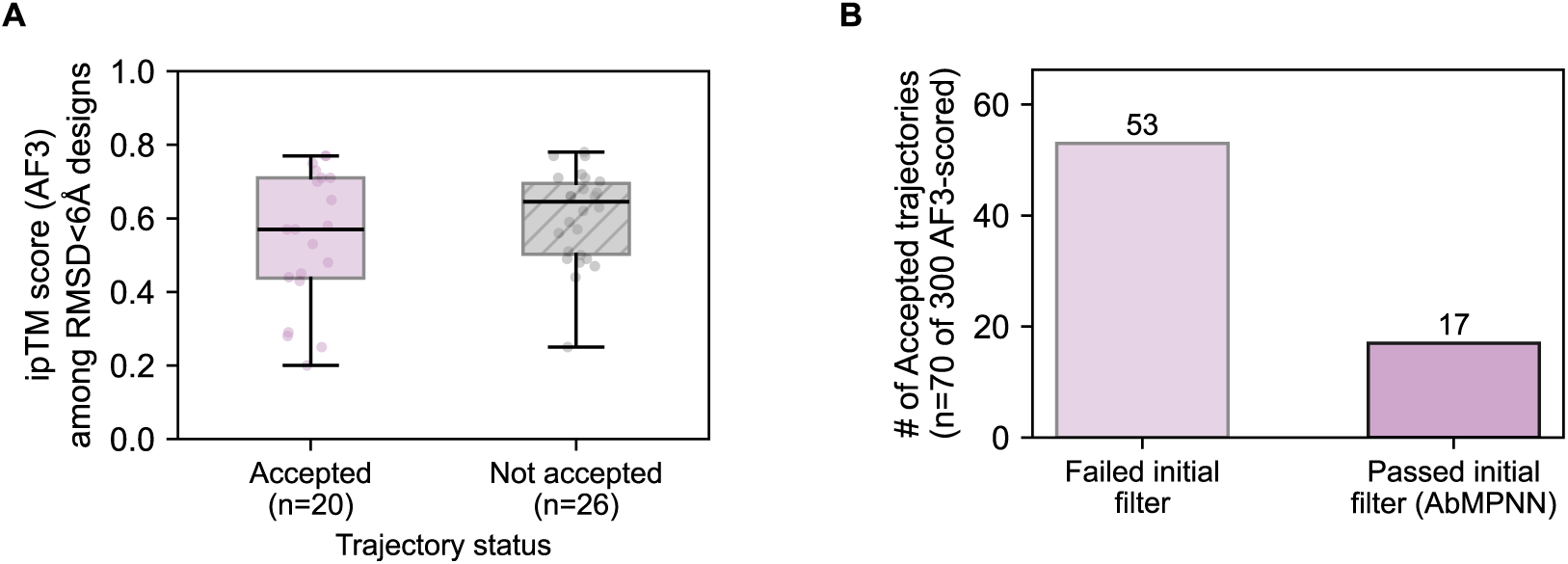
Germinal trajectory status and internal filtering: **A)** AF3 ipTM distribution among Germinal designs passing the RMSD self-consistency filter (RMSD *<* 6 Å), split by trajectory status: *Accepted* (Germinal accepted the trajectory) versus *Not accepted* (Germinal did not accept the trajectory), pooled across the three frameworks. **B)** Of the accepted trajectories (not filtered by AF3 or RMSD), the number that failed versus passed Germinal’s initial post-hallucination filter (Table 2) ahead of AbMPNN sequence redesign.

**Fig. S6.**
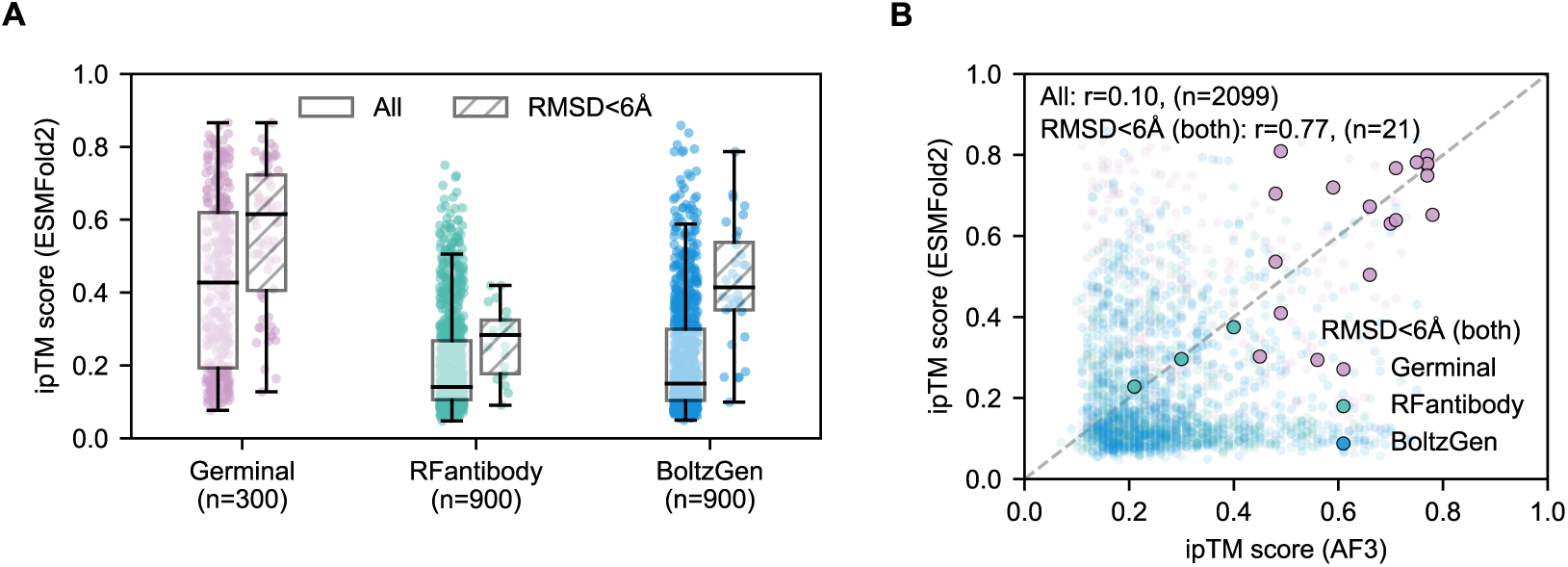
Model comparison cross-validation with ESMFold2: **A)** ipTM score per design method (Germinal, RFantibody, BoltzGen), re-scored with ESMFold2 instead of AF3, shown for all designs and for the RMSD *<* 6 Å self-consistent subset, reproduces the increased performance for Germinal for this target. **B)** Per-design ipTM agreement between AF3 and ESMFold2 on the same designs: essentially uncorrelated across the full population (Pearson r = 0.10, n = 2099), but restricting to designs both scorers call self-consistent (RMSD *<* 6 Å by both metrics) recovers a stronger correlation (r = 0.77, n = 21).

**Fig. S7.**
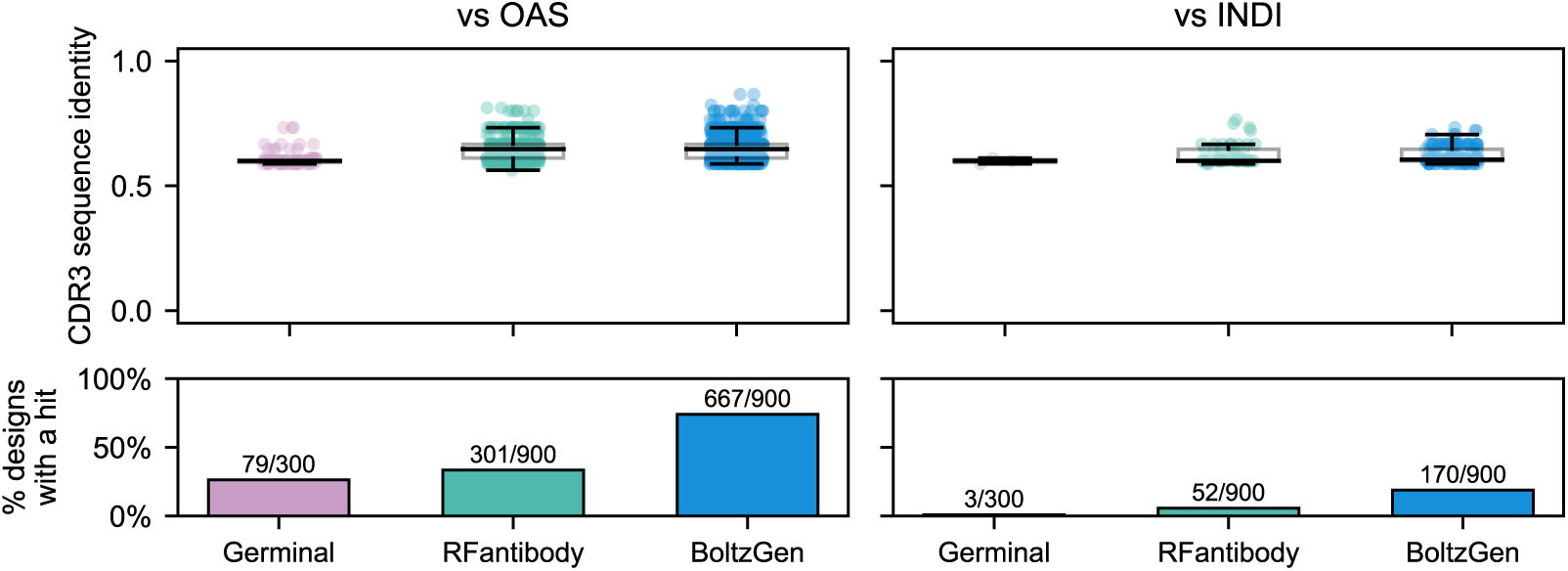
Sequence identity to antibody and V_H_H sequence databases: Generated V_H_H sequences from each design model were aligned against either the OAS (all antibody) or INDI (V_H_H-only) sequence databases. Unlike the SAbDab alignment in Fig. **1**D, MMseqs2 was run with a minimum sequence identity of 60% and minimum coverage of 90% to keep the search computationally tractable against these much larger databases. Top row: CDR3 sequence identity of the best hit, for designs that returned at least one hit above these thresholds. Bottom row: percentage of each model’s full design pool that returned any hit at all. Germinal produced the lowest hit percentage against both databases, consistent with Fig. **1**D.

**Fig. S8.**
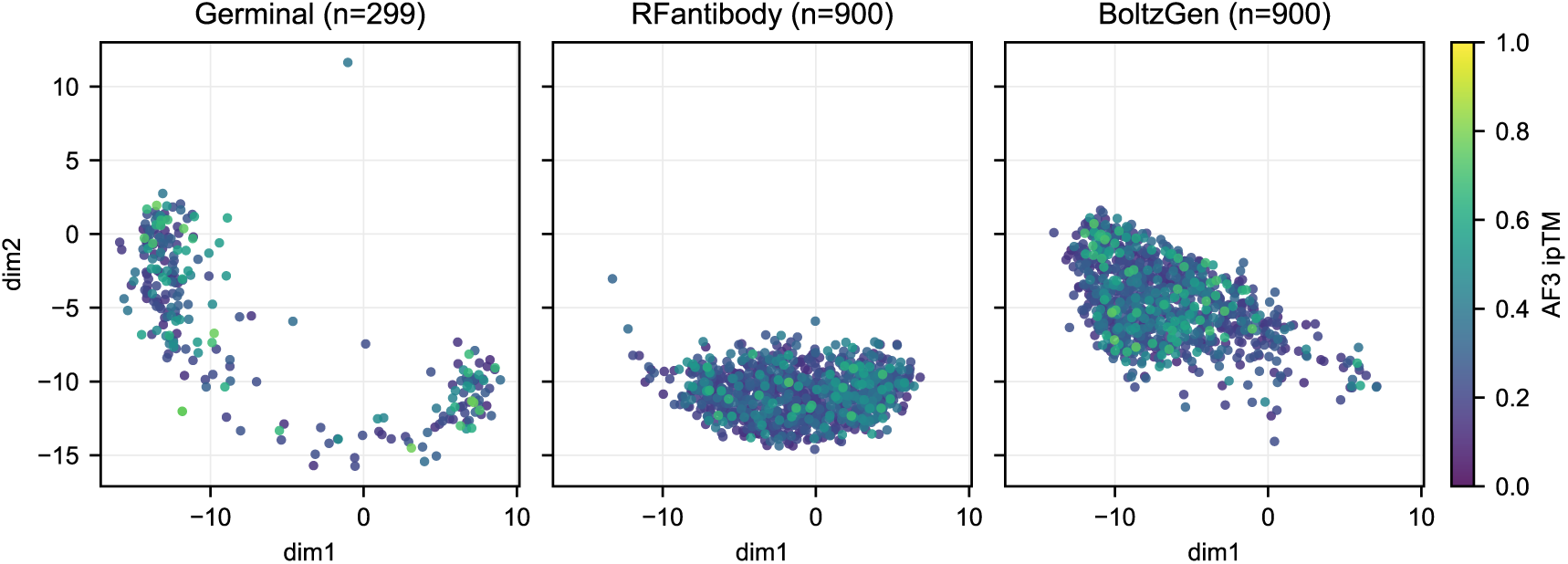
MDS projection by ipTM: The CDR3 MDS projection from Fig. **1**E, colored by AF3 ipTM and split by design method (background repertoire removed). Within each method’s own occupied region of CDR3 space, high- and low-ipTM designs are interspersed rather than clustered into a distinct sub-region. This indicates that high-confidence designs are not restricted to a defined region of CDR3 sequence space.

**Fig. S9.**
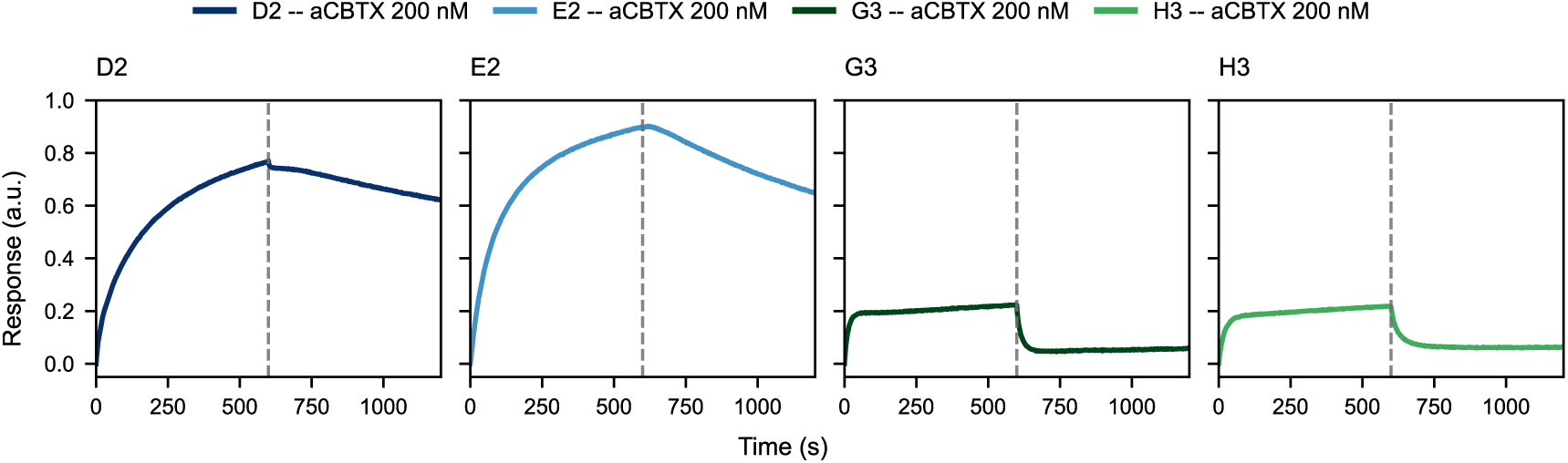
One-point BLI measurements of identified hit candidates: BLI response at a fixed concentration of 200 nM for D2, E2, G3, and H3. D2 and E2 show a substantially higher response than G3 and H3.

**Fig. S10.**
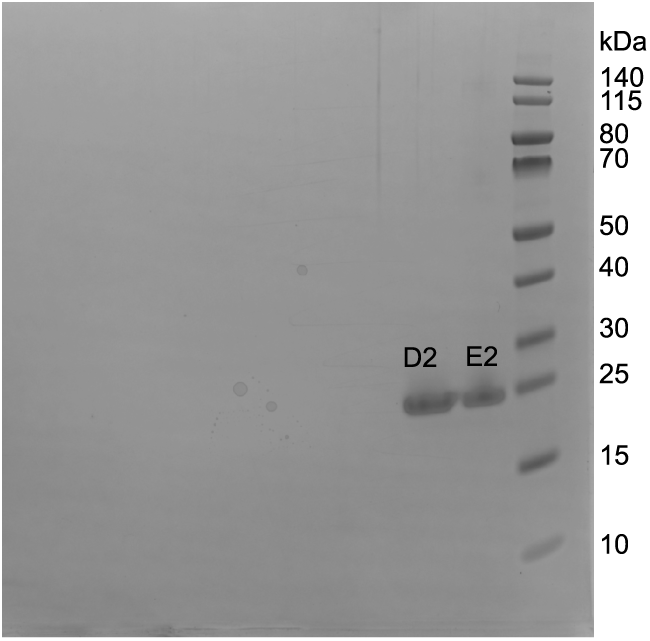
D2 and E2 purity. SDS-PAGE of D2 and E2 monomer fractions after SEC, confirming the expected molecular weight (*∼*18.7 kDa including tags) and high purity.

**Fig. S11.**
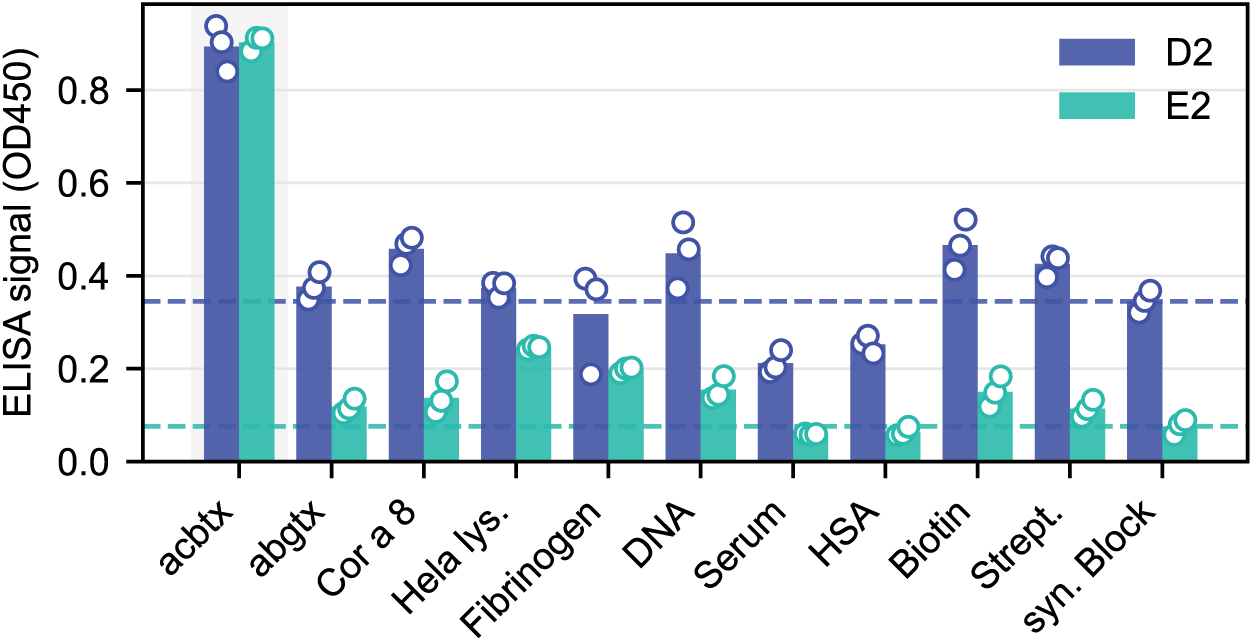
D2 and E2 polyreactivity at 100 nM: Polyreactivity at 100 nM V_H_H, showing increased signal against off-target antigens; however, background signal to the non-protein, non-DNA synthetic blocking-agent control (dashed line) was also elevated.

**Fig. S12.**
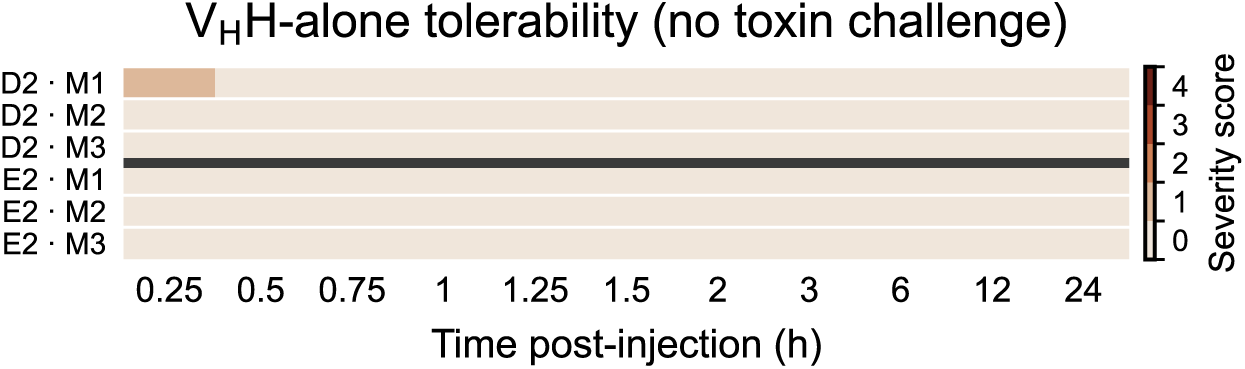
V_H_H-alone tolerability, per-animal severity heatmap: Clinical severity score for mice receiving D2 or E2 alone (no toxin/venom challenge), scored at 11 timepoints from 15 min to 24 h post-injection. Each row is one animal (*n* = 3 per cohort); color denotes severity score at that timepoint. No deaths occurred in either cohort.

## Supplementary Tables

**Table S1.** PDB structure presence in model training sets. Inclusion of the V_H_H frameworks and *α*-cobratoxin target structure (sourced from 9GCN) in the training datasets of each design method and structure predictor.

|  | 3EAK | 9GCN | 7XL0 |
| --- | --- | --- | --- |
| Germinal (AF2) | Y | N | N |
| RFantibody | Y | N | Y |
| BoltzGen | Y | N | Y |
| AF3 | Y | N | N |
| ESMFold2 | Y | N | N |

**Table S2.** Germinal V_H_H filtering criteria. Summary of the default initial (post-hallucination) filters and the more stringent final filters applied at the end of the Germinal pipeline. Dashes indicate criteria not applied at that stage.

| Filter metric | Initial filter | Final filter |
| --- | --- | --- |
| Steric clashes | < 1 | < 1 |
| Binder near hotspot | == **true** | == **true** |
| CDR3-hotspot contacts | > 0 | > 0 |
| Fraction of interface residues in CDRs | > 0.5 | $\geq 0.5$ |
| Interface shape complementarity | > 0.6 | $\geq 0.6$ |
| Self-consistency RMSD (sc_rmsd) | — | < 6.0 |
| Interface hydrogen bonds | — | $\geq 3$ |
| Surface hydrophobicity | — | $\leq 0.4$ |
| Interface hydrophobicity | — | $\geq 45$ |
| pDockQ2 | — | > 0.23 |
| External pLDDT | — | > 0.87 |
| External ipTM | — | > 0.80 |
| External pTM | — | > 0.80 |
| External PAE | — | < 7.5 |

**Table S3.**
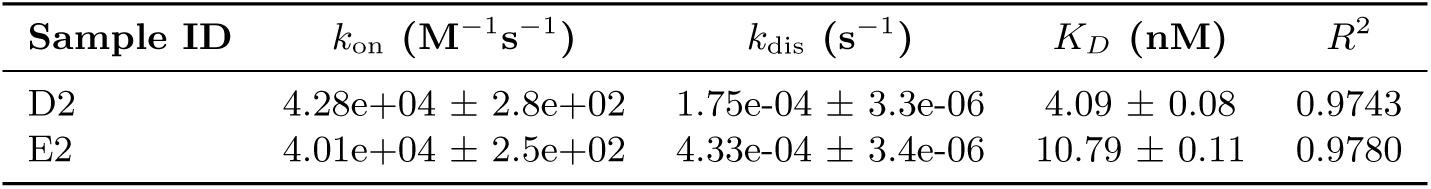
BLI kinetics for D2 and E2. Global 1:1 fit to a six-point, two-fold dilution series against *α*-cobratoxin.

